# Quantitative determination of fluorescence labeling implemented in cell cultures

**DOI:** 10.1101/2023.03.27.534369

**Authors:** Chiara Schirripa Spagnolo, Aldo Moscardini, Rosy Amodeo, Fabio Beltram, Stefano Luin

**Affiliations:** NEST Laboratory, Scuola Normale Superiore, piazza San Silvestro 12, I-56127, Pisa, Italy; NEST Laboratory, Istituto Nanoscienze-CNR, piazza San Silvestro 12, I-56127, Pisa, Italy

**Keywords:** Fluorescent labeling, Fluorescence microscopy, Single-molecule imaging, Single-particle tracking, Membrane receptors, Sfp phosphopantetheinyl transferase

## Abstract

**Background:** Labeling efficiency is a crucial parameter in fluorescence applications, especially when studying biomolecular interactions. Current approaches for estimating the yield of fluorescent labeling have critical drawbacks that usually lead them to be inaccurate or not quantitative.

**Results:** We present a method to quantify fluorescent-labeling efficiency that addresses the critical issues marring existing approaches. The method operates in the same conditions of the target experiments by exploiting a ratiometric evaluation with two fluorophores used in sequential reactions. We show the ability of the protocol to extract reliable quantification for different fluorescent probes, reagents concentrations, reaction timing and to optimize labeling performance. As paradigm, we consider the labeling of the membrane-receptor TrkA through 4’-phosphopantetheinyl transferase Sfp in living cells, visualizing the results by TIRF microscopy. This investigation allows us to find conditions for demanding single and multi-color single-molecule studies requiring high degrees of labeling.

**Conclusions:** The developed method allows the quantitative determination and the optimization of staining efficiency in any labeling strategy based on stable reactions.

## Background

Fluorescent labeling of cellular components is of utmost importance for studying biological processes since it enables the investigation of a variety of biological processes in sundry kinds of samples with high sensitivity, specificity and spatiotemporal resolutions [1, 2]. Advances in labeling strategies and microscopy yielded access to the single-molecule level and to the direct observation of biomolecule dynamics and interactions in physiological environments [3–8]. These homo- and hetero-interactions strongly correlate with molecular functions, as in the case of signal transduction via membrane receptors of different families, whose activity is mainly regulated by interactions with ligands, coreceptors and other membrane and intracellular molecules [9–13].

To obtain quantitative estimations of interaction parameters, it is essential to know the fluorescent-labeling efficiency, and this must be as high as possible [14–16]. The probability of observing an n-mer is proportional to the n^th^ power of the labeling fraction (assuming independency for labeling of different molecules) e.g., with a 20% labeling of dimerizing molecules, the probability of observing a dimer is only 4% [17]: this, even with relatively high -albeit incomplete-labeling efficiency, clearly indicates the need for corrections on experimental results. Yet several limitations affect existing approaches for labeling-efficiency estimation, as explained in the following.

Some studies report simple approaches based on the percentage of labeled cells or fluorescent-signal intensity to compare labeling strategies, tags or dyes [18–20], but these comparisons do not yield absolute efficiency values. Moreover, the measured parameters can include multiple effects, from variability in transfection efficiencies or expression levels [21, 22], to differences in dyes brightness or detection in different channels.

In other cases, labeling efficiency is evaluated via reaction kinetics studies on purified protein solutions [23–28], where conditions usually differ significantly compared to live cells, e.g. for protein accessibility, solubility and diffusion of the reactants, their permeability in cellular compartments, off-target reactions, toxicity. Moreover, protein purification may not be straightforward, especially for membrane proteins [29].

In other approaches, the labeling performance within a construct is estimated by searching for colocalizations between two fluorophores attached to a similar but different construct incorporating an additional fluorescent protein [30] or a second tag [31]. These methods demand the production and validation of additional constructs and moreover, experimental colocalization typically corresponds to proximity at distances of about 100-200 nm, due to optical resolution, while molecular-level colocalization implies nanometer-scale proximity [32–34]. In addition, especially for large probes such as Qdots or other nanoparticles, steric hindrance or other probe-probe interactions can interfere with the labeling of the same moiety with two probes [7, 35, 36]; in the case of fluorescent proteins included in the constructs, the fraction of non-fluorescent proteins (e.g. in metastable dark or immature states) can further influence the results [15].

Other authors test the efficiency by employing labeling with biotin followed by biotinylation analysis (e.g. Western blotting), even if the probe of interest is different [28, 37–39]. However, the efficiency may depend on the probe, as reported in some studies [24] and in the present work.

Here we present a robust ratiometric method to quantify the fluorescent-labeling efficiency of biomolecules, which overcomes the limitations described above. Importantly, it can be implemented in the same conditions of the target experiments, with the same sample, protein and probe of interest, by simply exploiting two sequential reactions with two different labels.

We show the power of the method in a complete procedure for labeling optimization, in which we identify the best compromise between high-specific and low-nonspecific dye interactions for different fluorophores. This compromise is an essential requirement to achieve challenging single-molecule applications [40–42].

We tested our method on TrkA receptors labeled by Sfp phosphopantetheinyl transferase [37, 39, 43]. In comparison to other approaches like self-labeling enzyme tags (like HALO or SNAP [7, 44]), this labeling strategy involves more reagents and therefore there are more parameters involved in its optimization; however, the measurement of labelling efficiency would be the same for these other approaches as well. The developed method allowed a thorough investigation of the efficiency varying all reaction parameters (concentrations, timing, fluorescent dyes) for the first time for any labeling reaction, to our knowledge. Moreover, we identified the conditions for single or multi-color single-molecule imaging at high degree of labeling, never reported before for Sfp-based labeling.

## Results

### Method to determine biomolecule-labeling efficiency

We targeted the quantification of labeling efficiency (i.e. the ratio between labeled molecules and the total number of accessible ones) by exploiting two sequential reactions, where the molecules available for the second reaction are the ones unlabeled in the first one. By using samples identical to those of interest, one can simultaneously assess the efficiency and the relevance of nonspecific interactions of the probes (often affecting fluorescent labeling [40–42]) in different conditions.

The workflow for the developed method is reported in Fig.1. A first labeling reaction is performed by attaching a fluorescent probe A, with efficiency *e*_*A*_; a second reaction with efficiency *e*_*B*_ is performed sequentially using a fluorescent probe B emitting in a different band. Considering a cell expressing N labelable molecules, there will be *e*_*A*_. *N* molecules labeled with A, leaving (*N* − *e*_*A*_*N*) labelable molecule for the second reaction; therefore, the molecules labelled with fluorophore B will be *e*_*B*_. (*N* − *e*_*A*_*N*). The ratio between the number of molecules labeled in the first and in the second reaction will be:

**Fig. 1.**
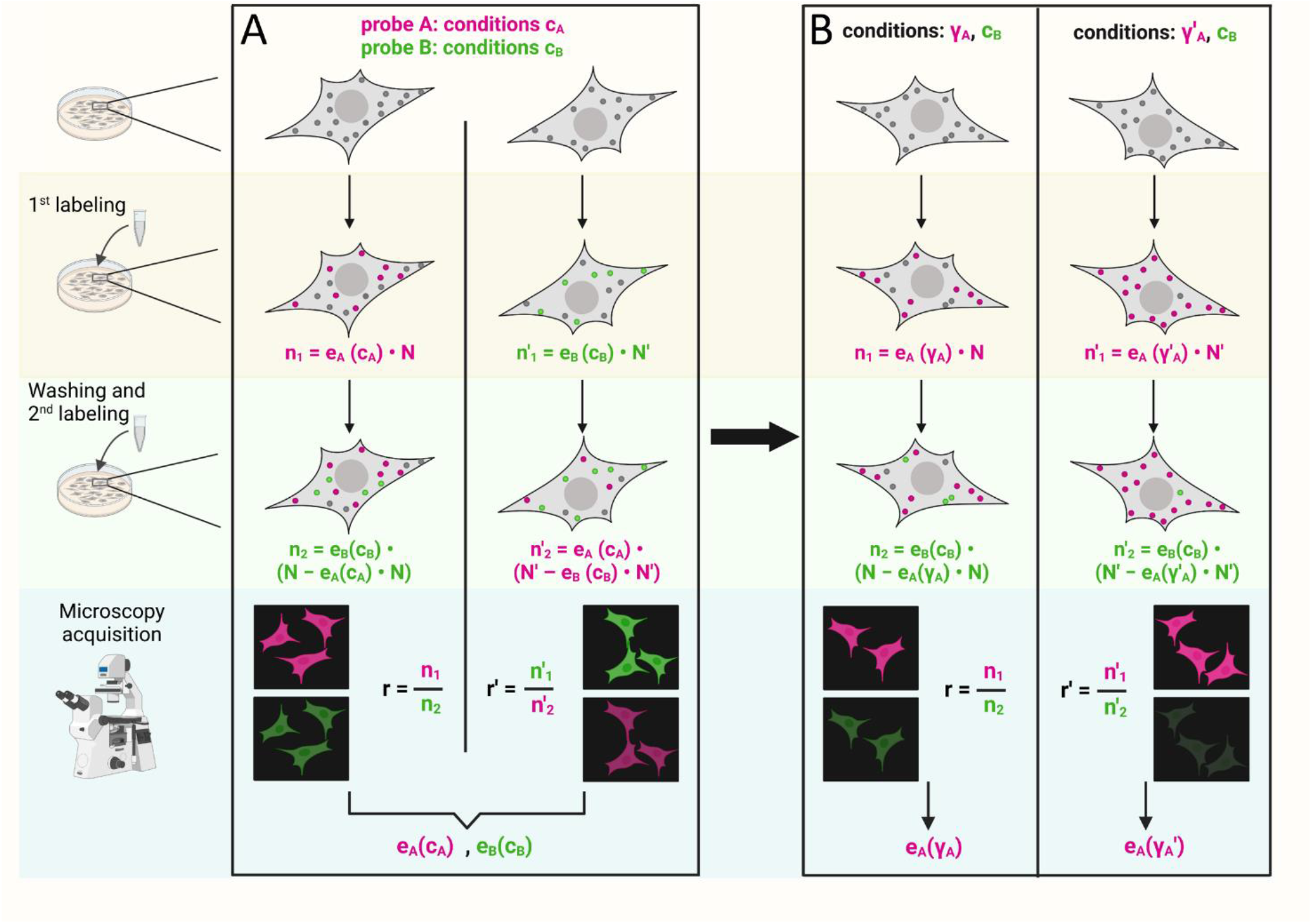
Method workflow. Left, outside boxes: depiction of the steps needed in the experiments. In panels A and B, each of the four columns represent a different experiment; grey circles in cells represent unlabeled molecules, which initially are N^(‘)^; magenta and green circles are molecules labeled with either of two different dyes (n^(‘)^_1_ or n^(‘)^_2_ in number, where subscript 1 or 2 indicates respectively the first or second labeling reaction). On the bottom, the two microscopy detection channels (in green and magenta) are schematically represented for a single field of view for each experiment to visualize the same cells labeled with both the two dyes at the end of the whole labeling procedure. Box A: method to be used when no labeling efficiency information is available; both experiments must be performed, inverting the order of labeling while keeping constant the reaction conditions c_A_ and c_B_ for the magenta and the green probe respectively. From the ratios of the labeled molecules r and r^’^ in the two cases, it is possible to extract the labeling efficiencies e_A_(c_A_) and e_B_(c_B_) for the two probes in the used conditions (see main text and equations (1) and (2)). Box B: method to measure e_A_ in varying conditions (e.g., γ_A_, γ_A_^’^) of the “magenta” dye, keeping constant the conditions c_B_ for the “green” one used as control with known efficiency e_B_(c_B_). Molecules are labeled sequentially with the test and the control probe and the ratio between the number of molecules labeled with the two dyes is evaluated. Each experiment yields the labeling efficiency e_A_(γ_A_), e_A_(γ_A_^’^) for the tested probe in the used condition (equation (3)). Created with BioRender.com

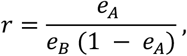

which depends only on the two labeling efficiencies and not on the expressed-molecule number *N* (see also Supplementary Note 1).

In a different sample, the two sequential labeling reactions are carried out in reverse order (Fig. 1A). The first reaction will label *e*_*B*_. *N* molecules; the second one will label *e*_*A*_ (*N* − *e*_*B*_*N*) molecules. The ratio between the number of molecules labeled in the first and in the second reaction will be:

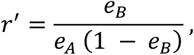

which again depends only on the two efficiencies.

The two equations can be solved simultaneously for *e*_*A*_ and *e*_*B*_:

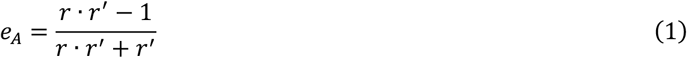

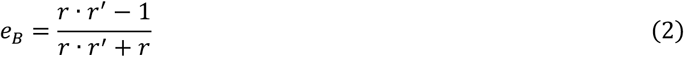

Starting from the measure of *r* and *r*^*′*^, the efficiencies for the two different probes in the used conditions can be determined. In order to investigate the reaction behavior as resulting from the variation of some parameters, one probe is selected as the probe of interest (“test probe”), with reactions at varying conditions, and the other as the “control probe”, with the previously-used labeling conditions at known efficiency (Fig. 1B). These experiments do not require performing the couples of reactions inverting the order. By labeling the cells first with the test probe (unknown efficiency *e*_*A*_) and then with the control probe (at known efficiency *e*_*B*_), the following relationship yields *e*_*A*_:

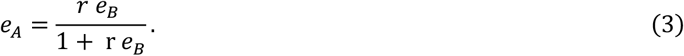

Otherwise, the control probe can be used in the first reaction (known efficiency *e*_*A*_) and the test probe in the second one (unknown efficiency *e*_*B*_) and the desired efficiency is:

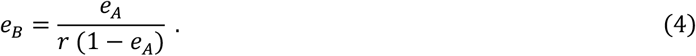

Usually, one measures a ratio of fluorescent intensities in two channels, therefore a relation between this ratio and the one of the number of labeled molecules must be determined or measured; in our case, we did this by measuring the single-molecule fluorescence intensity (see Methods). See also Supplementary Note 1 for a discussion of the application of similar, even if less precise, method using a single dye.

### Application to S6-TrkA fluorescent labeling through Sfp-phosphopantetheinyl-transferase varying dye and enzyme concentrations

We applied our method to characterize the fluorescent labeling of TrkA receptors on cell membrane through Sfp phosphopantetheinyl transferase (Sfp). In a Mg^2+^-dependent reaction, this enzyme covalently conjugates the phosphopantetheinyl arm of Coenzyme A (CoA) functionalized with any probe to a specific serine residue of a peptide tag inserted in the receptor sequence [37, 39, 45]. We employed the minimally invasive (12 residues) S6-tag, inserted in the TrkA sequence as previously shown [45–47] (Fig. 2A).

**Fig. 2.**
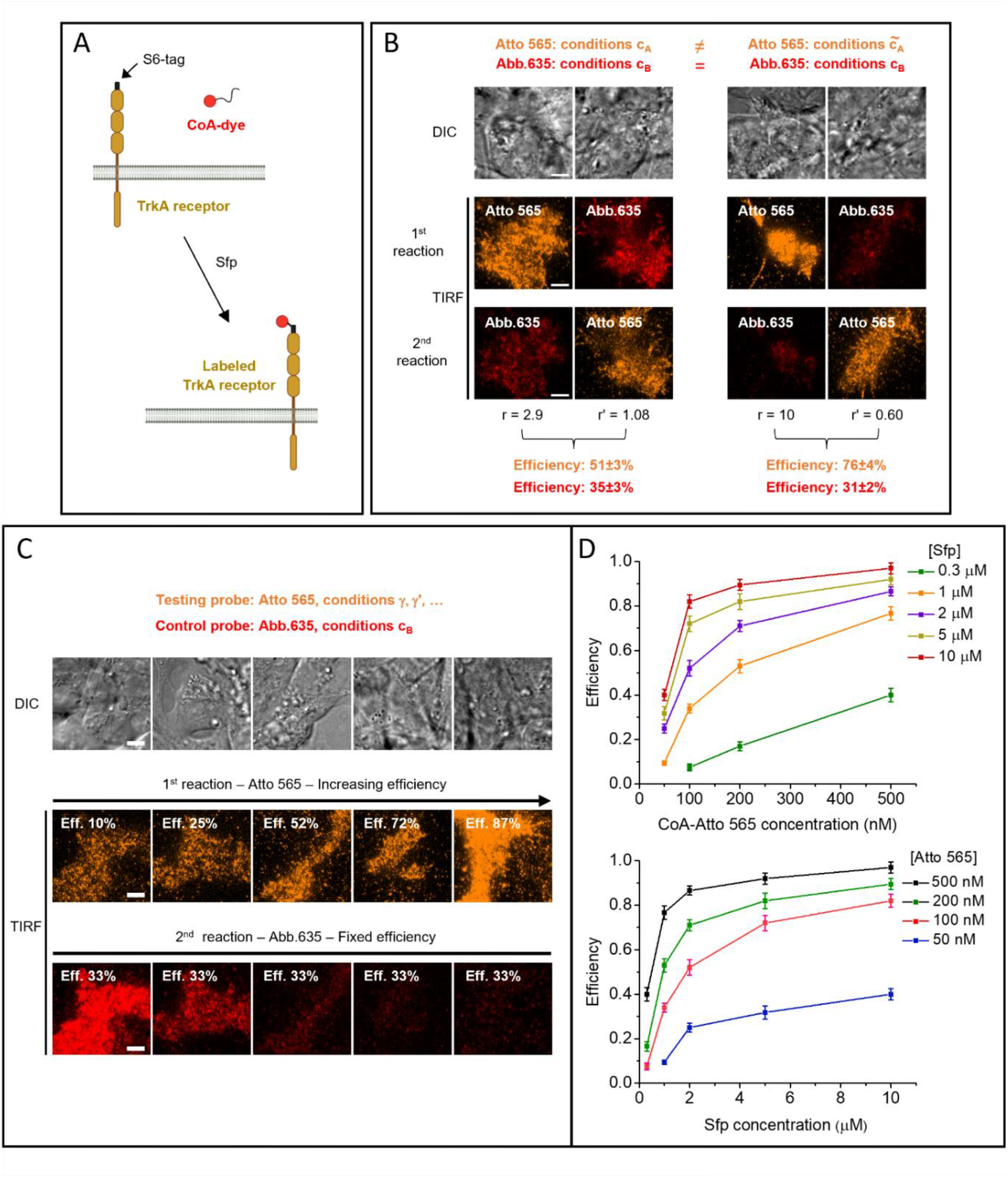
Application of the method to TrkA receptors labeled through Sfp. (A) Scheme for Sfp-based labeling applied on S6-tagged TrkA receptors expressed on living cell membranes. (B) Determination of labeling efficiency, performed as described in Fig. 1A, for Abberior STAR 635p (Abb.635, red channel) in conditions c_B_, while using two different conditions (c_A_ on the left and 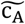 on the right) for reactions with the other probe (Atto 565, orange channel). Each image column shows a single representative cell: DIC image (top) and TIRF images of the channel of the dye used in the first (middle) and second (bottom) labeling reaction. Conditions for Atto 565 were: 100 nM CoA-dye, 2 µM Sfp and 20 min reaction time (left), and 100 nM CoA-dye, 10 µM Sfp and 10 min (right); conditions for Abb.635 were in both cases 40 nM CoA-dye, 10 µM Sfp, 20 min. r and r’ are the ratios between the number of molecules labeled in the two channels. Reported efficiency results are mean ± SEM estimate (see Methods), obtained from 17, 20, 25 and 16 cells analyzed for experiments represented in each image column. The results obtained for Abb.635 efficiency are not significantly different, and the efficiency estimated averaging the two results is 32.7±1.7%. (C) Examples of experimental images corresponding to the workflow steps depicted in Fig. 1B, for some of the analyzed conditions reported in panel D. Each image column shows a single cell: DIC image (top); TIRF channel of the tested probe Atto 565 (middle, orange), TIRF image of the control probe Abb.635 (bottom, red). Eff.: efficiency estimated (for Atto 565 in varying conditions) or previously measured (for Abb.635 in fixed conditions). (D) Fluorescent labeling efficiency obtained for Atto 565 at various concentrations of CoA-dye ([Atto 565]) and Sfp ([Sfp]), after 20 minutes of reaction and in presence of 10 mM of MgCl_2_ (dots: mean, error bars: SEM estimates; 15-40 cells from 2 independent replicates analyzed for each condition). Scale bar: 5 µm. See also Supplementary Tables 1 and 2

We studied the labeling efficiency with the organic dye Atto 565 (test probe), choosing the organic dye Abberior STAR 635p as the control probe. The ratios *r*^*′*^, *r* in equations 1-4 were determined from the total intensities of dyes inside labeled cells, knowing single molecule intensities and correcting for background and spuriously adsorbed fluorophores (see Methods).

First, we carried out the experiments involving the two labeling reactions in the two different sequential orders. As a first assessment for the robustness of control-probe efficiency estimation, we employed two distinct conditions for Atto 565 (Fig. 2B, Supplementary Table 1). Estimates for the control-probe efficiency were comparable.

Then we characterized the efficiency of Atto 565 in different situations.

Figures 2 C, D report the behavior at different concentrations of CoA-Atto 565 and Sfp, keeping fixed the reaction time at 20 minutes and the MgCl_2_ concentration at 10 mM. We demonstrate that our approach can explore the full range of labeling efficiencies, from a small percentage to complete labeling.

We checked the robustness of the method against control-probe conditions by performing experiments where we fixed the conditions for the test probe Atto 565 and we used two or three different control-probe situations, e.g. by varying control dye, its conditions or reaction order (Supplementary Fig. 1). Applying (3) or (4) with the corresponding value of the control probe efficiency (as previously determined, see Supplementary Tables 1, 2), we obtained comparable results for the tested probe efficiency, so the estimate proved stable and robust.

We compared our method with an approach based on simple receptor-density comparisons (Supplementary Fig. 2). As expected, the latter is too sensitive to protein expression variability and does not yield a reliable estimate of efficiency trends. On the contrary, our method demonstrated robustness against expression variability, thanks to the ratiometric approach (Supplementary Fig. 3).

We also monitored efficiency as a function of MgCl_2_ concentration by fixing the other reaction parameters (Supplementary Fig. 4): efficiency increases with MgCl_2_ concentration and saturates at 10 mM, the concentration typically used in this reaction [37]. We used this concentration in all the other experiments.

We checked if labeling efficiency is influenced by serum starvation before or during labeling, as typically performed in studying receptors stimulation (Supplementary Fig. 5), without finding significant differences.

### Labeling efficiency and non-specific interactions of the dye must be balanced for single-molecule imaging

As can be also seen in Fig. 2, the same labeling efficiency for a dye can be reached with different combinations of concentrations for the dye and for Sfp. However, the different conditions typically cause diverse amounts of dyes nonspecifically adsorbed to glass and cell membranes (“spurious adhesion”). This unwanted spots cause a background that can interfere with measurements, especially in single-molecule studies [40–42]. To investigate this, we measured the density of adhesion spots in areas including non-transfected cells after extensive washing following the reaction. We observed an increase of nonspecific spot density with increasing dye concentration; at 200 nM or above, the conditions are not compatible with single-molecule studies as the spot density becomes too high to distinguish individual molecules (Fig. 3A). Spurious adhesion also increased between 0.3 µM and 1 µM of Sfp (Fig. 2B), while there were no significant differences in the range 1-10 µM of Sfp. However, at 0.3 µM of Sfp, the efficiency was only 40% even with 500 nM of dye and the nonspecific density was already too high, therefore it is not possible to use this concentration of Sfp to achieve high efficiencies while limiting nonspecific spot density. Thus, the data achieved so far indicate that it is necessary to limit the dye concentration to 100 nM and to increase the Sfp concentration to 10 µM in order to reach high degrees of labeling (82%; see Fig. 2D) while limiting the density of spurious spots and allowing single-molecule studies.

**Fig. 3.**
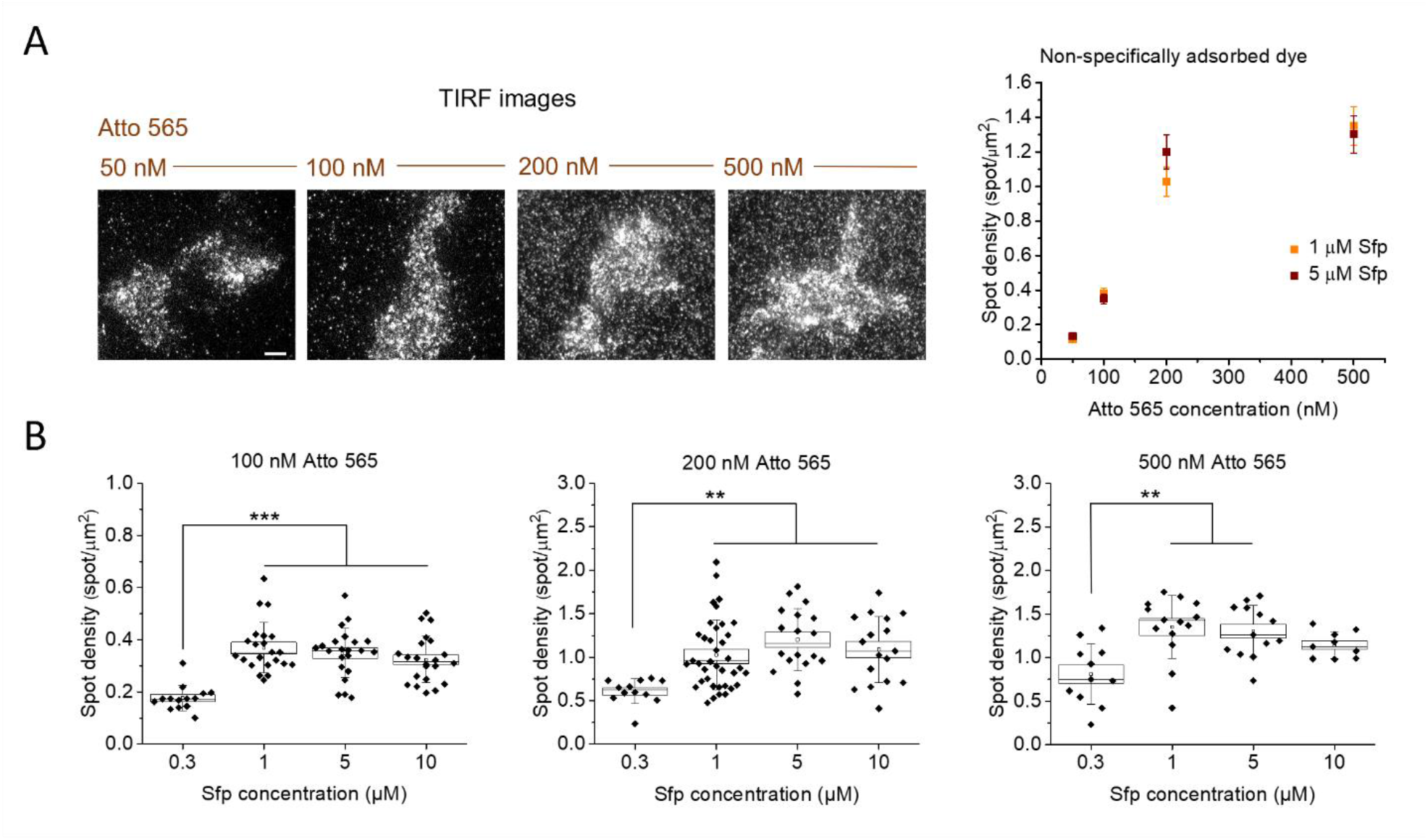
Non-specific adhesion of CoA-dye to cells and glass in the labeling reaction. (A) Left: representative TIRF images of cells incubated with a reaction mix including different CoA-Atto 565 concentrations (20-minutes reactions, 5 µM Sfp). Each field of view shows a single transfected labeled cell surrounded by dyes non-specifically adsorbed to glass or other non-transfected cells (see Methods). Scale bar: 5 µm. Right: quantification of spot density for non-specifically adsorbed dyes using different CoA-Atto 565 concentrations in the reaction, at 1 and 5 µM of Sfp (mean ± SEM from 12-38 fields of view analyzed in two independent replicates). Spot density is measured after 20 minutes of reaction and extensive sample washing (see Methods). (B) Non-specific dye adhesion varying Sfp concentration, at different fixed CoA-Atto 565 concentrations (box: mean ± SEM, whiskers: standard deviation, empty circles: averages, horizontal lines: medians, diamonds: individual data from two independent replicates). ***P < 0.001, **P < 0.01, 1-way Anova, Bonferroni multiple comparisons; no significant difference if nothing is shown

Results in Figs. 2 and 3 were obtained using a fixed incubation time of 20 minutes. We then investigated the behavior of specific and nonspecific dye interactions at different incubation times for the labeling reaction (Fig. 4 A, B; Supplementary Table 2). We used concentrations of 100 and 200 nM for Atto 565, each one with 1 and 10 µM of Sfp. The main point in this characterization concerns the individuation of the efficiency value at plateau and the needed time to reach it. For both, we observed differences between the analyzed conditions. Increasing enzyme concentration increases significantly the plateau efficiency value, even if this is reached in a longer time, while increasing dye concentration increases both plateau efficiency and reaction rate.

**Fig. 4.**
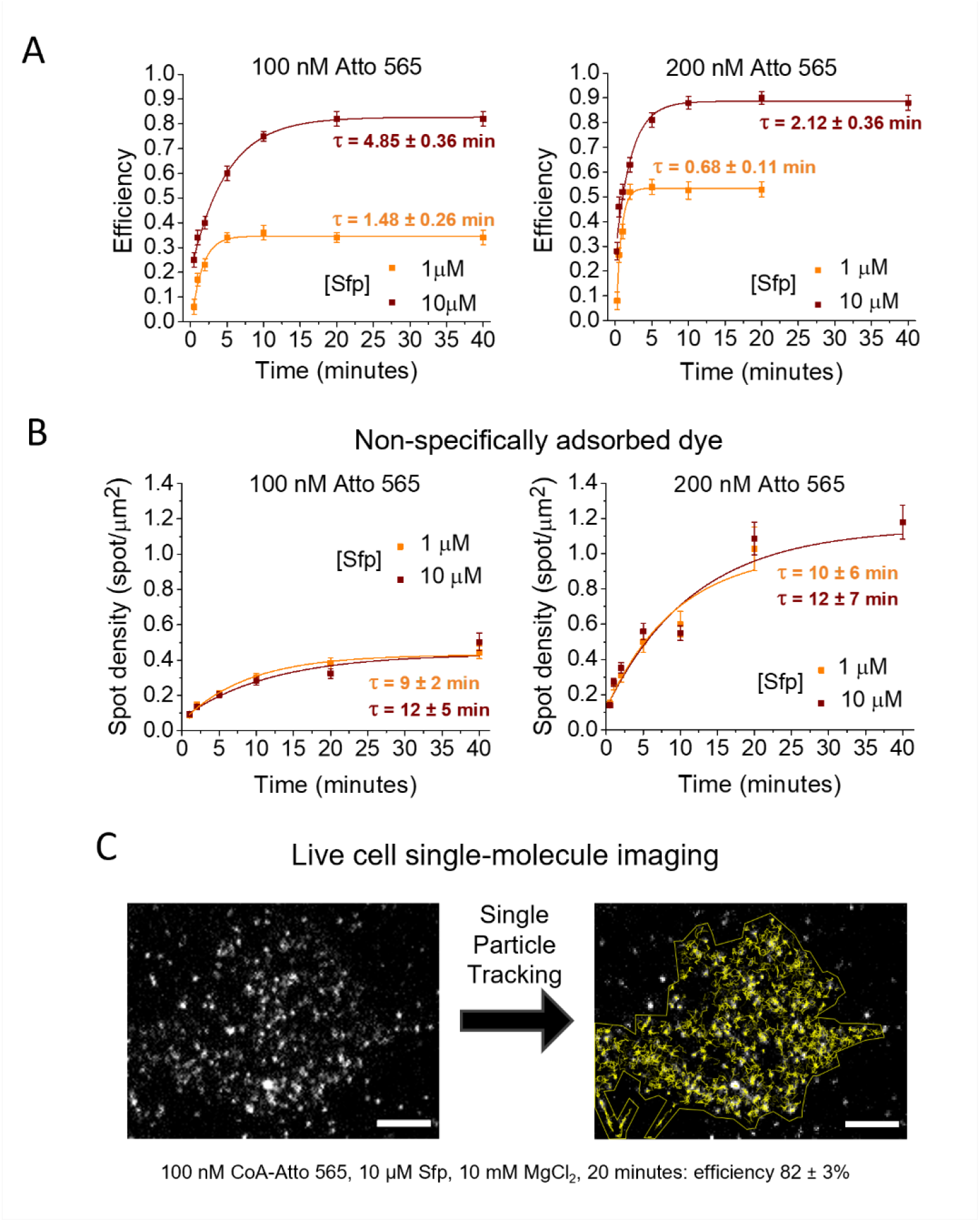
Labeling efficiency and nonspecific adhesion as functions of time. Conditions for single-molecule imaging at high labeling efficiency. (A) Labeling efficiency as a function of time measured at 1 (orange) and 10 (bordeaux) µM of Sfp at CoA-Atto 565 concentration of 100 nM (left) and 200 nM (right). (B) Density of spots non-specifically adsorbed to cells and glass as a function of time measured in the same conditions of panel A. Dots are means, error bars are SEM estimates, continuous lines are mono-exponential fitting curves (with obtained lifetime τ in minutes). 15-30 cells from 2 independent replicates were analyzed for each condition. (C) Single-molecule imaging in live cells and application of single-particle tracking analysis at found optimal labeling conditions (100 nM CoA-Atto 565, 10 µM Sfp, 10 mM MgCl_2_, 20 minutes: efficiency 82 ± 3%). A representative cell is shown. Gray scale: TIRF images; left: first frame of the acquired 100-frame movie; right: in yellow, tracks obtained from a single-particle tracking analysis on frames 1-50, superimposed on the 50^th^ frame (see also Supplementary Videos 1, 2). Scale bars: 5 µm

In Fig. 4B we also report the behavior of nonspecific adsorption as a function of time; most significantly, nonspecific adsorption still increases even when efficiency reaches a plateau in every studied condition. These observations highlight the requirement of using the shortest possible incubation time to obtain the wanted efficiency level: further increasing time would only increase the density of spurious adhered spots. More considerations on optimum dye concentration could arise concerning the combinations of concentrations and reaction times for reaching the best compromise between high efficiency and low spurious adhesion. Indeed, while the results of Figs. 2D and 3 (obtained with reaction times of 20 minutes) suggest limiting dye concentration to 100 nM, one should consider that the time required for completing the reaction decreases with increasing dye concentration and that nonspecific density decrease for shorter reaction times. From the plots in Fig. 4 A, B, it is possible to directly compare efficiency and spurious density reached at a dye concentration of 100 nM after 20 minutes with the same quantities reached at 200 nM after 10 minutes (in both cases with 10 µM of Sfp and with plateau efficiency practically reached). Spurious density is higher in the second case, with only a minor increase in efficiency. Moreover, at the time when a dye concentration of 200 nM produces a nonspecific density comparable to that observed under the conditions identified as potentially the best at 100 nM (i.e. ∼0.35 spots/µm^2^ at around 2 minutes for 200 nM dye concentration), the efficiency is lower. Therefore, we can confirm 100 nM as the optimal dye concentration, but the considerations reported above highlight the necessity of simultaneously evaluating specific and nonspecific dye interactions.

Using the established reaction conditions (100 nM CoA-Atto 565, 10 µM Sfp, 10 mM MgCl_2_, 20 minutes), we performed fluorescent labeling in live cells expressing the s6-TrkA receptor, demonstrating a single-particle tracking example at high labeling efficiency in this system (Fig. 4C and Supplementary Videos 1, 2).

### Different fluorescent dyes require different conditions in the reaction

We compared the behavior of different fluorescent dyes in the labeling reaction (Fig. 5A). Atto 565 and Abberior STAR 635p showed comparable efficiencies in the analyzed reaction conditions (Fig. 5B and Supplementary Table 1). On the contrary, three 488-dyes (Alexa, Atto, Abberior STAR) showed lower efficiency in comparison to Atto 565 (Fig. 5C). We did more in-depth comparisons between Atto 565 and Atto 488 (Fig. 5D-F, Supplementary Table 2). We measured efficiencies at different dye concentrations (from 50 nM to 500 nM), keeping fixed Sfp at 10 µM and time at 20 minutes: in order to obtain similar efficiencies for the two fluorophores, about twice the concentration is required for Atto 488 (Fig. 5E). We compared the efficiency of Atto 488 at 200 nM and Atto 565 at 100 nM as a function of time and we observed the same kinetics for the two dyes, with an efficiency of Atto 488 around 90% the Atto 565 one at each time. Fig. 5F shows significant differences also for the nonspecific adsorption of the two dyes under equal conditions. As is the case for efficiencies, these spurious adhesions are comparable for 200 nM of Atto 488 and 100 nM of Atto 565 at each reaction time.

**Fig. 5.**
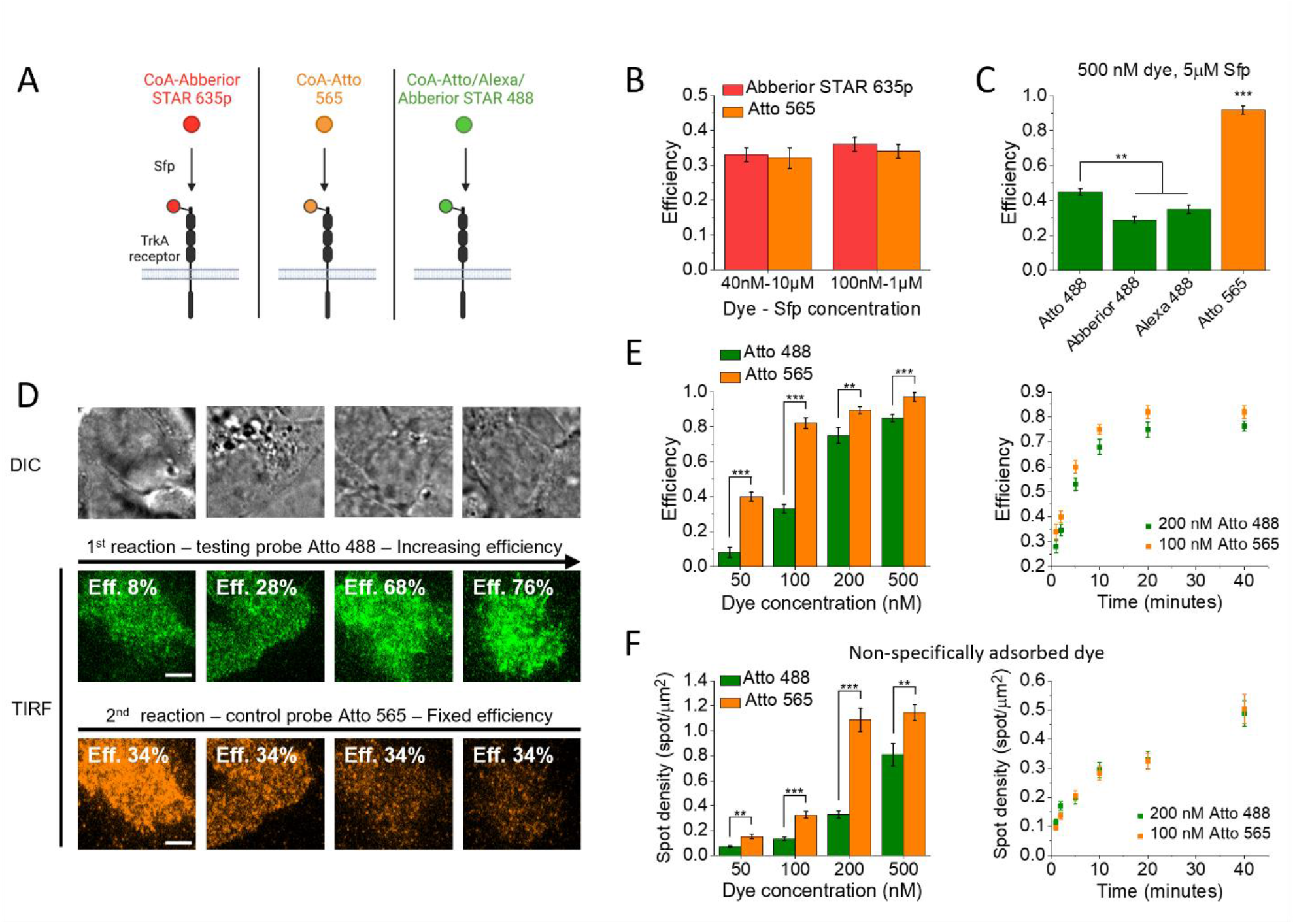
Comparison of different fluorescent dyes in the labeling reaction. (A) Sketch of labeling with different fluorescent dyes, emitting in wavelength bands represented by the used colors. Created with BioRender.com. (B) Labeling efficiency of Atto 565 and Abberior STAR 635p at two different couples of dyes-Sfp concentrations (reaction time: 20 minutes). (C) Efficiency for Atto 565 compared with Atto 488, Alexa 488 and Abberior STAR 488 for 20-minute reactions in fixed reaction conditions. ***P < 0.001, **P < 0.01, 1-way Anova, Bonferroni multiple comparisons. (D) Experiments for measurement of Atto 488 efficiency in variable conditions. The first labeling is performed with the test probe Atto 488, the second labeling is performed with the control probe Atto 565, the latter in conditions of known efficiency. Each image column shows a single cell: DIC image (top), TIRF image of Atto 488 channel (middle, green), TIRF image of Atto 565 channel (bottom, orange). Eff.: estimated efficiency. Scale bar: 5 µm. (E) Efficiency comparison between Atto 488 and Atto 565. Left: comparison at different dye concentrations ([Sfp]: 10 µM, time: 20 minutes). ***P < 0.001, **P < 0.01, Welch test. Right: comparison at different reaction times at the indicated dyes concentrations ([Sfp]: 10 µM). (F) Comparison of non-specifically adsorbed spot density for Atto 488 and Atto 565, using the same conditions as in panel E. Results on Atto 565, already shown in previous figures 2, 3, 4 are here reported for direct comparison. ***P < 0.001, **P < 0.01, Welch test. In B, C, E, F data are mean ± SEM and are obtained from 13-30 analyzed cells from two independent replicates.

There is some evidence in the literature on correlation between dye hydrophobicity and spurious interactions, but these same studies report various exceptions to this behavior and the involvement of multiple factors [40–42]. Anyway, authors detected a lower level of nonspecific interaction for Atto 488 compared to Atto 565, as we obtain here for similar concentrations in the reaction. A lower hydrophobicity is reported for Alexa 488, Atto 488 and Abberior STAR 488 compared to Atto 565 and Abberior STAR 635p [19, 40–42]; thus, hydrophobicity could be an important factor also for labeling efficiency, at least in the system we studied.

These results highlight the significance of our method for dye comparison because it allows the direct evaluation of the probe of interest. Moreover, they confirm the need to inspect simultaneously labeling efficiency and nonspecific interactions since both can differ for different probes. Even if Atto 488 requires higher concentrations compared to Atto 565 to reach the same labeling efficiency, its lower level of nonspecific interactions makes these higher concentrations more usable.

### Two-color balanced labeling of TrkA receptors

We investigated the conditions for simultaneous two-color labeling of the TrkA receptor. Labeling a protein simultaneously in different channels is a powerful approach for studying homodimerization processes through techniques like multicolor single-particle tracking or FRET [48–51]. In this case, there is the additional requirement of a specific ratio between the efficiency of different dyes, used simultaneously in the labeling but typically requiring different conditions, just like in the cases we examined. In two-color single-particle tracking, it is desirable to have a ratio close to one between the two channels for efficient detection of homodimerization events (see Supplementary note 3). We performed simultaneous two-color labeling of the TrkA receptors using CoA-Atto 565 and CoA-Atto 488 mixed in the same reaction. Because of the different efficiency of the two dyes, we used a twofold excess of Atto 488. To evaluate the labeling efficiency, the mix Atto 565 + Atto 488 was the “test probe” for our method, while we used Abberior STAR 635p as control probe (Fig. 6 A, B and Supplementary Tables 1, 2). We tested concentrations that, based on the previous results, were expected to correspond to high-efficiency values (Fig. 6C). Besides the total efficiency of the Atto 565 + Atto 488 mix, we also evaluated the ratio between the receptors labeled with each of the two dyes. Interestingly, at 100 nM Atto 565 + 200 nM Atto 488 with 10 µM of Sfp, we observed a lower relative efficiency of Atto 488 (with a 488-to-565 ratio of 0.6) while with 20 (or 30) µM of Sfp we obtained a ratio of 0.9, comparable to the one shown in Fig. 5E. The plateau efficiency of 100 nM Atto 565 + 200 nM Atto 488, 20 µM of Sfp was equivalent to that of 200 nM 565, 10 µM Sfp (in both cases about 90%), even if in the two-color case it was reached at a shorter time. The conditions 100 nM Atto 565 + 200 nM Atto 488, 20 µM Sfp, 10 mM MgCl_2_ and 5 minutes reaction time were the optimal choice to allow two-color single-molecule experiments at high labeling efficiency, with balanced labeling between the two channels and a tolerable amount of nonspecific adhesion. In Fig. 6D and Supplementary Videos 3, 4, 5 we show an example of two-color single-particle tracking performed in these conditions.

**Fig. 6.**
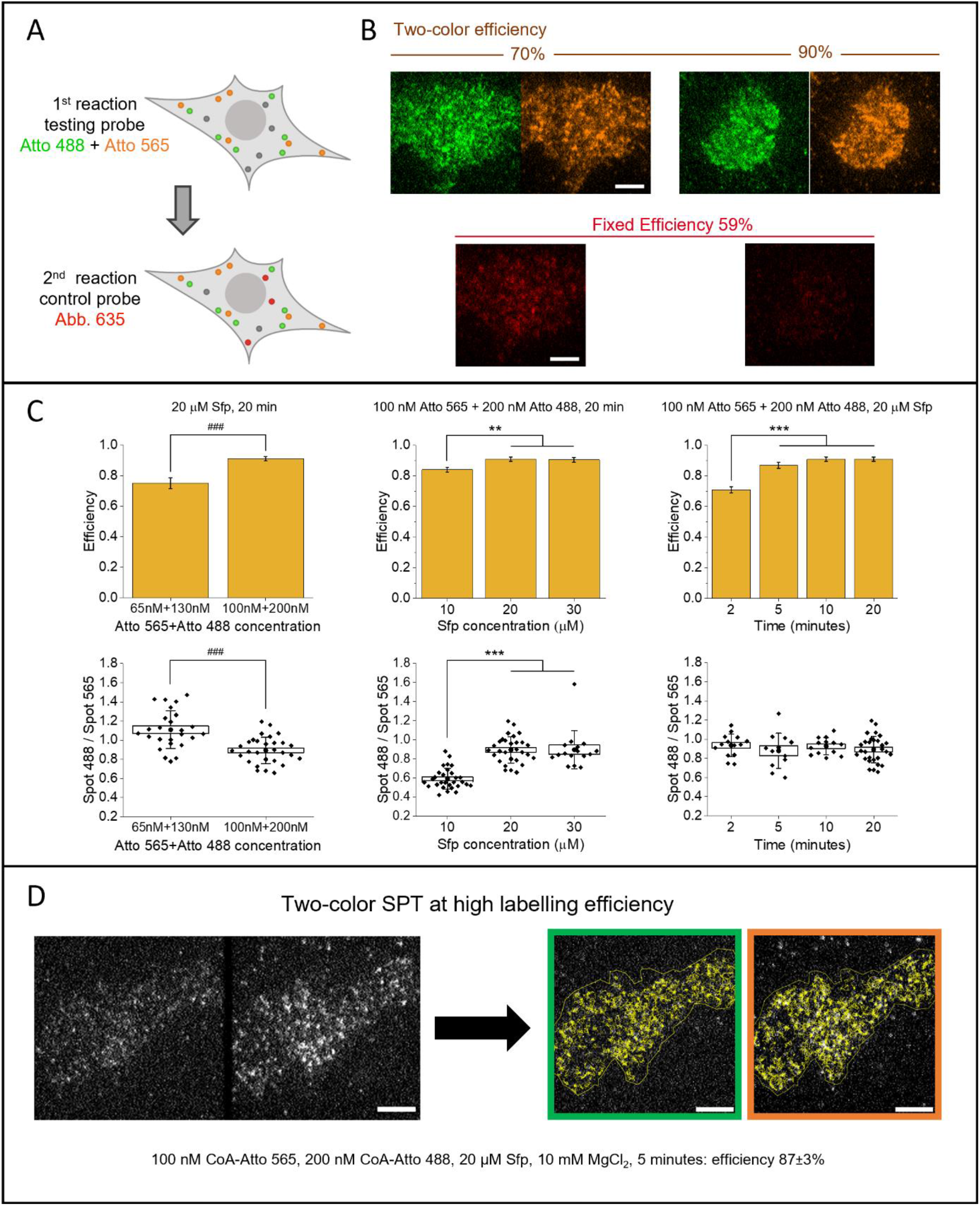
Conditions for simultaneous two-color labeling. (A) Sketch of the experiment for efficiency measurement in simultaneous two-color labeling. The test probe is a mix of Atto 565 and Atto 488; the control probe is Abberior STAR 635p. Created with BioRender.com. (B) Experimental TIRF images acquired after the two labeling reactions. For each of the two representative shown cells, we reported at the top the Atto 488 (green) and Atto 565 (orange) channels, with estimated total efficiency for the two-color labeling; at the bottom the Abberior STAR 635p channel (red); scale bar: 5 µm. (C) Results in different reaction conditions as described above the graphs for each column; top: measurement of two-color labeling efficiency (mean ± SEM), bottom: ratio between the number of molecules labeled with Atto 488 and with Atto 565 (box: mean ± SEM, whiskers: standard deviation, dots: averages, diamonds: individual data from two independent repetitions). ***P < 0.001, **P < 0.01, 1-way Anova, Bonferroni multiple comparisons. ^###^P < 0.001, Welch test. (D) Example of simultaneous two-color single-molecule imaging and single-particle tracking analysis at high labeling efficiency in the optimized conditions. Left: first TIRF image of the acquired movie where the same cell is simultaneously detected in the Atto 488 channel (left) and the Atto 565 channel (right). Right: superimposed yellow tracks as obtained with single-particle tracking after splitting and analyzing each channel (shown is the 50^th^ frame with tracks reconstructed until that frame from a 100-frame movie; see also Supplementary Videos 3, 4, 5). Scale bars: 5 µm

## Discussion and Conclusion

We introduced a method to quantify the fluorescence-labeling efficiency of biomolecules significantly improving over existing protocols. We tested its ability and robustness to measure efficiencies covering the full percentages range. We showed its reliability and the importance of its operating in the same conditions required by the target experiment by demonstrating: i) the different performance of different dyes; ii) the simultaneous investigation of specific and nonspecific dye interactions to achieve conditions for challenging single-molecule studies at high labeling efficiencies. Our method also applies to the characterization of two-color labeling performed on one molecule, so it is a powerful tool for studying homodimerization phenomena. The extension to this case is very straightforward, while there are no similar examples in the literature. The developed protocol does not require any additional expertise, equipment or reagents over the ones required for the experiment of interest, except for one additional fluorophore and determining the correspondence between the intensity ratio in the two channels and the ratio between the number of fluorescent molecules; this is very beneficial for minimizing cost and labor. Moreover, in Supplementary Note 1 we describe a similar method that, while less precise and not easily applicable in every kind of experiment especially with living cell, does not have these requirements.

Application of our quantitative labeling efficiency measuring protocol on TrkA receptor labeled by Sfp phosphopantetheinyl transferase shed new light on this reaction in the context of interest for fluorescence-microscopy applications. This labeling system offers the advantages of short and minimally-invasive tags and the versatility to attach a variety of probes. However, there was no characterization of its efficiency in this context, most likely due to the lack of a suitable method. The efficiency of this labeling strategy was reported by conjugating CoA-biotin in a fixed condition and examining the biotinylation [38]: a reaction performed with 50 µM of CoA-biotin, 0.5 µM of Sfp, 10 mM MgCl_2_ for 30 minutes yielded to a biotinylation of more than 80%. However, we observed a very high level of background due to spurious adhesion of dyes already at fluorophore concentration 100-times lower, around 500 nM (Fig. 3A), demonstrating that the proposed concentration of CoA-biotin [38] is not suitable for CoA-dyes on cells, especially for single-molecule studies.

Existing protocols for this labeling strategy report CoA-substrate concentrations ranging from few nM to tens of µM, with a typical enzyme concentration of 1-2 µM and with incubation times variable from 20-60 minutes to overnight [37, 52]. The highest substrate concentrations (order of 1-10 µM) are typically used for CoA-biotin, CoA-dyes on purified proteins, cells with very high expression levels, or in confocal microscopy applications; these are all cases where the background signal has less impact [37, 53–56]. Our results confirm that such concentrations (with the typical Sfp concentration in the low micromolar range) correspond to high labeling efficiency; however they cannot be used for single-molecule microscopy in cells. Indeed, in reported studies of this kind, CoA-probe concentration is reduced to the order of 1-10 nM because the application is sensible to dye molecules non-specifically adherent to coverglass or cell membrane [57–59]. Such low concentration values are suitable to study biomolecule-diffusion dynamics, but, in our case, we observed efficiencies not high enough to investigate biomolecule-interaction mechanisms.

Sfp-labeling is not the only system challenged by nonspecific adhesion of fluorophores. This is a problem encountered in many labeling strategies; e.g. self-labeling enzyme tags, like HALO, SNAP and CLIP tag [7, 40, 42, 60–62]; in general, nonspecific adsorption of dyes and proteins is a complex phenomenon affecting rather diverse bio-applications, from single-molecule imaging to biosensor development [63–65]. Optimizing rigorously the labeling parameters, as shown here, could establish the optimal conditions for efficient labeling limiting aspecific adhesion; in principle, having more parameters to tweak should help this optimization, but good results could be obtainable also for simpler labelling approaches, like the cited self-labeling enzyme tags where the only parameters are dye concentration and reaction time. The simultaneous check of efficiency and spurious adhesion is crucial also when comparing different probes. In the literature, there are some studies where the second factor alone is used as a criterion to exclude or select dyes [40, 42, 60]. These conclusions neglect that dyes considered better for lower levels of nonspecific interactions could require higher concentrations to get the same labeling efficiency of others with higher spurious effects, as in our comparison between Atto 488 and Atto 565.

The developed method can be applied to a variety of probes and biomolecules and to all labeling approaches based on a reaction realizing a stable probe conjugation on a system with a defined number of binding sites. It is not limited to labeling in cells, but it can be used in other kinds of systems, e.g. in tissues or on immobilized purified molecules. New probes and labeling strategies are continuously developed to investigate new processes, obtain better resolution, or be less invasive. Our method can be a true enabler of these new approaches for the study of biomolecule interactions, which are crucial mechanisms for their function. Additionally, as in the case of labeling by Sfp, it opens new perspectives for established techniques, not yet optimized for single molecule studies at high labeling efficiency, pushing further their applicability to gain further and more detailed insight into biomolecular behavior.

## Methods

### Synthesis of CoA-fluorophores

Alexa 488 maleimide was obtained from Molecular Probes (Invitrogen, Eugene, OR); Atto 488 maleimide and Atto 565 maleimide were obtained from ATTO-TEC (Siegen, Germany); Abberior STAR 635P maleimide and Abberior STAR 488 maleimide were obtained from Abberior (Goettingen, Germany). All solvents were purchased from Sigma-Aldrich and were of ultrapure grade. Coenzyme A trilithium salt was also purchased from Sigma-Aldrich. Chromatographic analyses and purifications were performed using a Phenomenex Kinetex EVO C-18 150 × 3.00 mm column on a Dionex Ultimate 3000 high-performance liquid chromatography system (HPLC) equipped with a photo diode array (PDA) detector. Eluent A: Ammonium acetate buffer 10 mM (pH 7.0). Eluent B: Acetonitrile:Eluent A (95:5). For the analyses, the HPLC was interfaced with an ABSciex API 3200 Q-Trap mass spectrometer (MS). MS parameters: Curtain gas 25.0 mL/min; Ion spray voltage 5500 V; Temperature: 100 K; Declustering potential: 40.0 V; Entrance potential: 10.0 V; Collision energy: 10.0 eV. For the purifications, the same HPLC was used, interfaced with a fraction collector. The quantifications of the CoA-dyes were performed with a UV/vis spectrophotometer (Jasco V-550). All CoA-fluorophore couplings were performed in the same way. CoA (0.412 µmol) was previously dissolved in a Tris(2-carboxyethyl)phosphine (TCEP) solution (10 mM in PBS, 100 µL) and stirred at 37 °C for 1 h. The solution was then cooled to room temperature and the fluorophore (0.375 µmol) dissolved in DMF (40 µL) was added. The reaction mixture was stirred in a thermomixer at 37°C for 1-3 h using times previously optimized checking when there was no further decrease of the starting reagent (fluorophore) by controlling a small quantity of the ongoing reaction mixture by HPLC-MS. The reaction mixture was purified by RP-HPLC. Finally, the CoA-dye was lyophilized and stored at -20°C in the dark. Before use, it was reconstituted in PBS and the concentration was determined through absorbance measurements at the dye’s maximum using published extinction coefficients.

### Purification of 4’-phosphopantetheinyl transferase Sfp

Production and purification of 4’-phosphopantetheinyl transferase Sfp was performed according to the protocol in [57] with minor changes. Briefly, after we centrifuged the bacterial suspension expressing the DNA plasmid coding for the PPTase enzyme at 6000 g, 4 °C for 20 min, we resuspended the obtained pellet in 10 mL of lysis buffer composed by 20 mM Tris-HCl at ph=8, 300 mM NaCl, 30 mM imidazole supplemented with protease inhibitor tablets (cOmplete™, EDTA-free Protease Inhibitor Cocktail, Sigma Aldrich), 0.05% of TritonX and 1 µg/ml of lysozyme. This suspension was sonicated on ice with six pulses of 30 s, separated by 60-s pauses. We then spun down the solution by two subsequent centrifugations at 13,000 g and 4 °C for 30 min and finally filtered the cleared lysate through a 0.45-μm syringe filter. For the purification step, we followed exactly the protocol described for the purification of His-tagged proteins using a gravity-flow column (HisPur™ Ni-NTA Resin (88221)-Thermo Fisher Scientific). Finally, the purified fraction was aliquoted and stored at -80°C in a solution of 20 mM Tris-Hcl at pH=7.5, 150 mM NaCl and 25% of glycerol.

### Construct

S6-TrkA construct was obtained by cloning the human cDNA of S6-TrkA described previously [45] into a pcDNA3.1+ vector. Briefly, starting from the “all-in-one” inducible lentiviral vector of the S6-TrkA construct, the full-length cDNA was amplified with PCR using FW (CGCCCAAGCTTACGCGTATGCTGCGAGGCGGACGG) and RV primers (CGATCTAGACGCACGCGTTCAGCCCAGGACATCCAGGTA) and then inserted between the XbaI and HindIII restriction sites of the pcDNA3.1+ vector.

### Cell culture, s6-TrkA receptor expression and labeling

SH-SY5Y cells (ECACC 94030304, a kind gift from Fondazione EBRI, Rome, Italy) were grown at 37°C, 5% CO_2_, in DMEM/F-12 medium supplemented with 10% Fetal Bovine Serum, 1% Penicillin-Streptomycin, 1% L-Glutamine. Two or three days before the microscopy experiment, cells were seeded in WillCo-dish® Glass Bottom dishes; at about 70% of confluency, they were transfected with the S6-tagged human TrkA by means of Lipofectamine 2000 (Thermo Fisher Scientific) according to the manufacturer instructions. After 24-36 hours, we performed the labeling reactions. Reactions were carried out in the culture medium (unless differently stated) using a mix containing Sfp Synthase, MgCl_2_ and the CoA-conjugated form of the organic dye. Reagent’s concentrations were varied as stated for the different experiments. Cells were incubated with the labeling mix at 37°C, 5% CO_2_ for a time varied as stated for the different experiments. In the case of two sequential labeling reactions, after the first one, we performed three washes with warm PBS and immediately incubated the cell with the reaction mix for the second labeling. See also Supplementary Tables 1 and 2 for details on concentration reagents, reaction times, and eventually order of reactions. At the end of the entire labeling procedure, cells were washed five times with warm PBS. In the case of live cells imaging, they were immediately imaged under the TIRF microscope; in the case of fixed cells, they were fixed for 90 min at room temperature with 4% PFA/2% Sucrose solution supplemented with 0.1% Glutaraldehyde (GA, Electron Microscopy Sciences), washed five times with PBS, kept in PBS and then imaged at the TIRF microscope. For two-color single-particle tracking measurements, 1mM Trolox (Sigma-Aldrich) and 5µM n-propyl gallate (Sigma-Aldrich) are included in the imaging medium to limit photobleaching effects over time [8].

### Experiments for determinations of efficiency in simultaneous two-color labeling with Atto 565 and Atto 488

For evaluation of simultaneous two-color labeling efficiency with Atto 565 and Atto 488, we prepared a reaction solution mixing the two dyes (with the concentrations stated in the different tests) with Sfp and MgCl_2_. After washing, in a following second reaction, we used as control probe Abberior STAR 635p at conditions: 70 nM CoA-dye, 10 µM Sfp, 10mM MgCl_2_, 20 minutes, corresponding to a control efficiency of 59%.

### TIRF microscopy

Cell imaging was performed using a Leica DM6000 inverted microscope (Leica Microsystems) equipped with an epifluorescence module, DIC in transmission, TIRF-AM module, HCX PL APO 100X oil-immersion objective (NA 1.47), electron-multiplying charge-coupled-device (CCD) camera (iXon Ultra 897, Andor), four laser lines (405 nm, 488 nm, 561 nm, 635 nm) and incubator chamber to maintain 37 °C and 5% CO_2_ conditions for live-cell imaging. Camera parameters were chosen to obtain the best compromise between signal-to-noise ratio and temporal resolution: the temperature was set to -75° C, pixel clocking rate to 17000 MHz, vertical shift speed to 0.5 µs, and vertical clock voltage to +2.

In experiments on cells labeled with both Abberior STAR 635P and Atto 565, each cell was imaged in two different channels using the 635 nm laser line coupled with a Cy5 filter cube and, separately, the 565 nm laser line coupled with a Cy3 filter cube. Powers at the objective were typically 3.5 mW and 3 mW for 561 nm and 635 nm laser respectively. In measurements on cells labeled with Atto 565 and one 488-dye, the two channels were detected simultaneously using a Quad filter cube and a Dual View (Optical Insights DV-CC, with filters Chroma ET525-50 and Semrock 600/52 nm BrightLine® and dichroic beam splitter 565dcxr) placed in front of the camera after an Optomask adjustable field mask (OPTMSK-L, Andor) [6, 8]. The Dual View windows in the different channels were aligned using bright field imaging of the supplied grid, leaving some unilluminated pixels between the two detection windows.

Simultaneous double excitation was achieved with the internal laser line at 565 nm plus an external 488 nm laser (iFLEX-iRIS, Qioptiq) connected through a laser combiner (iFLEX-adder, QiOptiq), using kineFLEX polarization-maintaining fibers (QiOptiq) and kineMATIX fiber couplers (QiOptiq) [6, 8]. Typically, laser powers at the objective were 3.2 mW and 3.5 mW for 565 nm and 488 nm lasers, respectively. The power of the external 488 nm laser was tuned using a DAC channel from a Daq2000 board. In measurements on cells labeled with the three dyes Abberior STAR 635p, Atto 565 and Atto 488, for each cell, two setups were alternated for two different time series: the first was the one designated for detection of the two channels 565 and 488 described above, the second was based on 635 nm excitation, obtained using the internal laser line passing through the laser combiner (with a typical power at the objective of 1.5 mW), coupled with the Cy5 filter cube. Typically, for each channel of interest, a 100-frame time series was acquired with a 40-ms integration time, resulting in a frame time of 56 ms.

### Determination of the number of labeled receptors in the cell basal membrane

To obtain the ratios of labeled receptors between different channels required in equations 1-4, we determined the number of labeled receptors *N*_*λ*_ for the dye *λ* in the observed cell basal membrane. We exploited the following quantities: the integral intensity detected within the observed basal membrane *I*_*in*_, the integral intensity of single-molecule spots *I*_*spot*_, the background and/or spurious signals, all determined as better explained below and in Supplementary Fig. 6. Our estimation automatically corrects for the different brightness and signal-to-background ratio of the different fluorescent dyes and can be applied to cells with various expression levels, from single-molecule densities to bulk ones. All quantities were obtained using ImageJ software.

Experiments were carried out on living cells, with transfected cells being identified by the movement of labeled receptors. First, we selected some transfected cells with labeled receptors density allowing us to distinguish single moving spots (Supplementary Fig. 6A). For each cell, we used the maximum intensity projection of the TIRF movie to identify the ROI of the observed portion of the basal membrane of transfected cells (cell ROI). Exploiting the TrackMate plug-in [66], we performed single-particle detection and tracking within this cell ROI. We optimized the spot radius parameter required by the software during the detection step and then used it to estimate the area of the spot *area*_*spot*_, assuming it to be circular (note that the spot area corresponds to the 2D point spread function as recorded by the CCD, which depends only on the microscopy setup). This quantity was then kept constant for all cells. From the tracking step, we obtained the integral intensity of each single spot included in tracks, as provided by the software. We selected tracks lasting at least 4 frames to exclude false detections and applied filters on displacement and velocity of tracks to exclude the immobile spots adhered to glass or membrane. We mediated the integral intensities of all resulting spots for each selected cell. We evaluated the background intensity *I*_*backgr*_ in the same movies, selecting a region of area *area*_*backgr*_ without spots neither in movement nor immobile (Background ROI), calculating a mean intensity through the whole movie and extracting the background intensity for unit area. We repeated these operations for different cells and the final quantities *I*_*spot*_ and 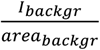 were calculated by taking the average over all the cells and the imaged fields. Of note, *I*_*spot*_ must be determined under conditions that exclude the detection of receptor oligomers or clusters to reliably estimate the intensities actually caused by single molecules. Unstimulated TrkA observed at very low density satisfies this requirement [58].

The theoretical brightness of the different fluorescent dyes we employed differs, and their distinct detection channels can have varying background levels. We obtained a relationship of direct proportionality between 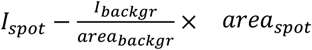 (i.e the background-corrected single-spot intensity) and the theoretical brightness multiplied by the experimental laser power in different setups for different dyes (Supplementary Fig. 6B). This confirms the reliability of our estimations. We then analyzed cells with arbitrary density of moving spots (Supplementary Fig. 6C). We again identified each cell ROI, having area *area*_*in*_, as explained above, and measured the integral intensity *I*_*in*_ inside *area*_*in*_ ; it includes the signal from the dyes labeling the receptors, the dyes non-specifically adsorbed to glass or cell membrane and the background originated by different sources such as autofluorescence of media, glasses or cells. In the same field of view of the transfected cell, we considered a region (of area *area*_*out*_) outside that cell; it was outside the transfected cell but included other non-transfected cells, because we used samples with cells at full confluency, as observed in DIC mode. In this area, we evaluated the integral intensity *I*_*out*_, which includes all the contributions that are in *I*_*in*_, except for the signal from the dyes labeling the receptors. To avoid photobleaching effects, *I*_*in*_ and *I*_*out*_ were computed from the first image in the movie.

Finally, we determined the number of labeled receptors *N*_*λ*_ using the following expression for each detection channel and each cell:

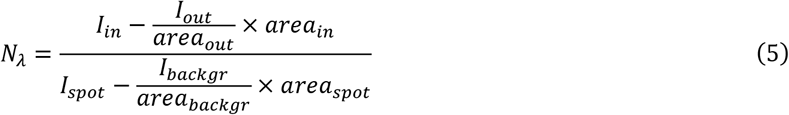

### Analysis of non-specific adhesion

We analyzed nonspecific adhesion density as shown in Supplementary Fig. 7, by selecting regions outside the transfected cells. These regions included non-transfected cells thanks to cell confluence; in this way, we could include both the adhesion to the glass substrate and to the cell membrane. We averaged the first 10 frames of the TIRF movie to improve the signal-to-noise ratio (SNR) of the single spots and to avoid having a significant number of missed detections for dyes with lower SNR, as in the case in Supplementary Fig. 7, where Atto 488 has a relatively low SNR in the raw image in comparison to Atto 565. On this averaged image, we used the detection step of the TrackMate plugin to obtain the spot density around the transfected cell.

### Comparisons of labeled receptor density

We compared the densities of labeled receptors using the same dye (Atto 565) in different conditions so that the density is simply proportional to the observed signal intensity. We carried out experiments both in fixed and living cells (Supplementary Fig. 2). We identified cell contours using the maximum intensity projection of TIRF movies in live cells and TIRF and DIC images in fixed cells. We extracted the mean intensity inside the transfected cell and subtracted the mean intensity outside the transfected cell by using ImageJ software.

### Statistics

Uncertainties on the efficiencies calculated using equations (1-4) were calculated by propagating the uncertainties on the quantities appearing on the right-hand side of those equations, considering that these were obtained from independent measurements (see Supplementary note 2). As uncertainties on averaged quantities, we consider their associated standard errors (SEMs).

The effective degrees of freedom that have to be used for comparison statistical tests have been calculated via the Welch-Satterthwaite approximation [67, 68], starting from the standard deviations and the (effective) degrees of freedom (*v*_*i*_ = *n*_*i*_ − 1 for a quantity averaged over *n*_*i*_ data) of the quantities entering in their calculation (see Supplementary note 2).

In all statistical tests, for p-values above 0.05 the differences were not marked as significant. If the p-value was under 0.05, we indicated the minimum significance among 0.05, 0.01 and 0.001. Used tests and sample sizes are stated in figure legends.

## Acknowledgments

We thank Laura Marchetti for useful discussions and manuscript revision. We acknowledge her and Fulvio Bonsignore’s work in developing the initial versions of some of the constructs used in this work.

## Authors’ contributions

Conceptualization, C.S.S. and S.L.; Methodology, C.S.S., S.L., R.A., A.M.; Validation, C.S.S.; Software, C.S.S.; Formal Analysis, C.S.S.; Investigation, C.S.S.; Resources, F.B., S.L.; Data Curation, C.S.S.; Writing – Original Draft, C.S.S., S.L.; Writing – Review & Editing, C.S.S., R.A., A.M., F.B., S.L.; Visualization, C.S.S., S.L.; Project Administration, S.L.; Supervision, S.L., F.B.; Funding Acquisition, F.B., S.L.

## Availability of data

The dataset generated during the current study are available from the corresponding author on reasonable request.

## Declarations

