## Supplementary Information for "Quantitative determination of fluorescence labeling implemented in cell cultures"

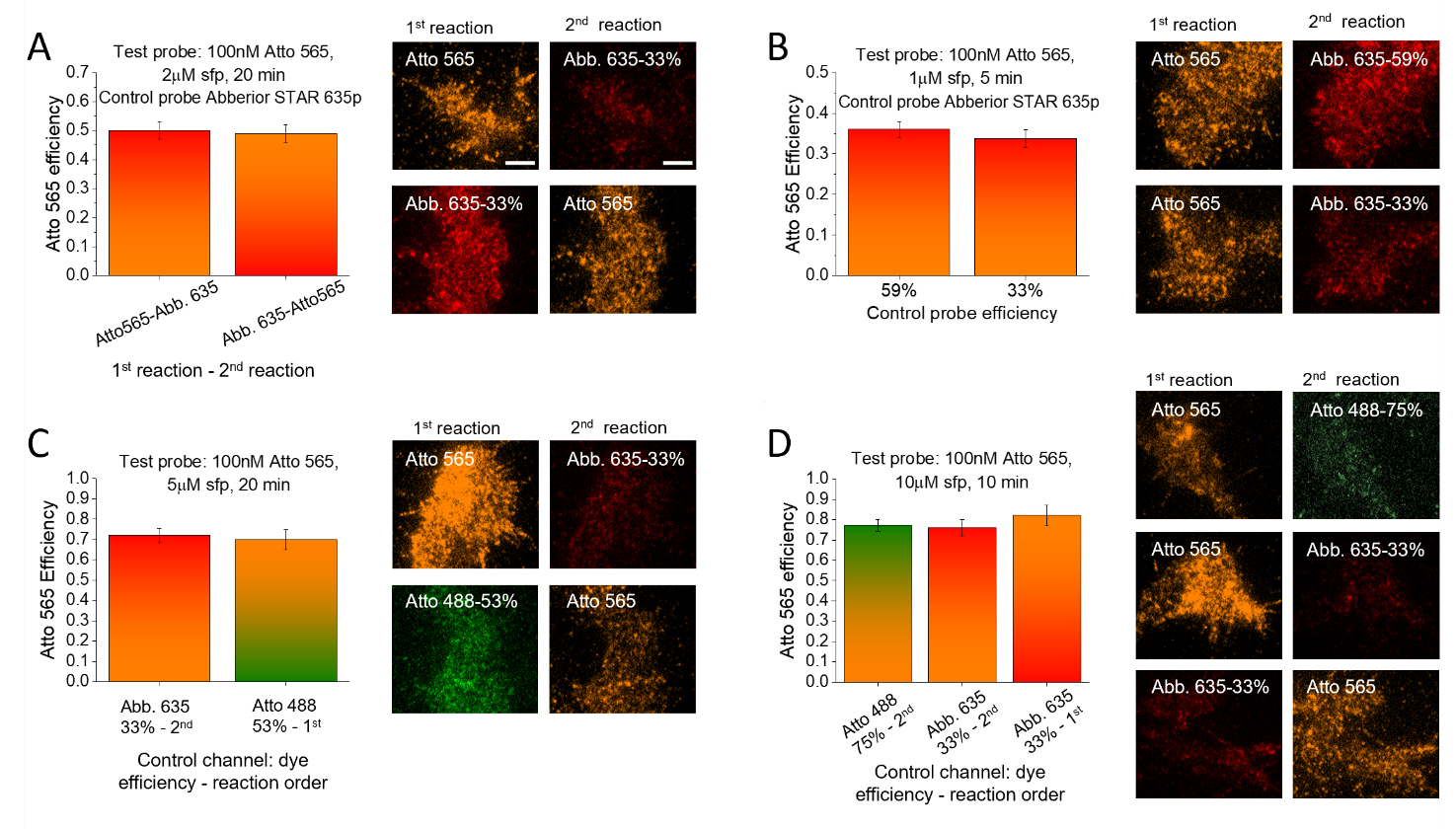

**Supplementary Fig. 1** Robustness of the efficiency estimation against control channel conditions. Atto 565 is the test probe at the conditions indicated within each panel, while two or three different situations are adopted for the control channel. See Supplementary Table 2 for control probe conditions and for the experiments where their efficiencies were measured. Bar plots show efficiencies estimations for Atto 565, reported as mean ± SEM (determined as described in Methods and in Supplementary Note 2; 10-22 cells from two independent replicates were analyzed in each condition). Bars colors indicate the two performed reactions: colors represent dyes (orange for Atto 565, red for Abberior STAR 635p, green for Atto 488) and gradients represent reactions order (bottom: first reaction, top: second reaction). On the right of each plot, representative TIRF images of the two acquired channels are shown (scale bar: 5 μm). (A) Control reaction with Abberior STAR 635p (Abb. 635) is carried out as the second or the first reaction. (B) Control reaction with Abb. 635 is carried out in conditions corresponding to two different control efficiencies. (C) Two different dyes with two different efficiencies and different reaction orders are used as control probes. (D) Control reactions characterized by different probe, efficiency, reaction order. Differences are not significant in any panel.

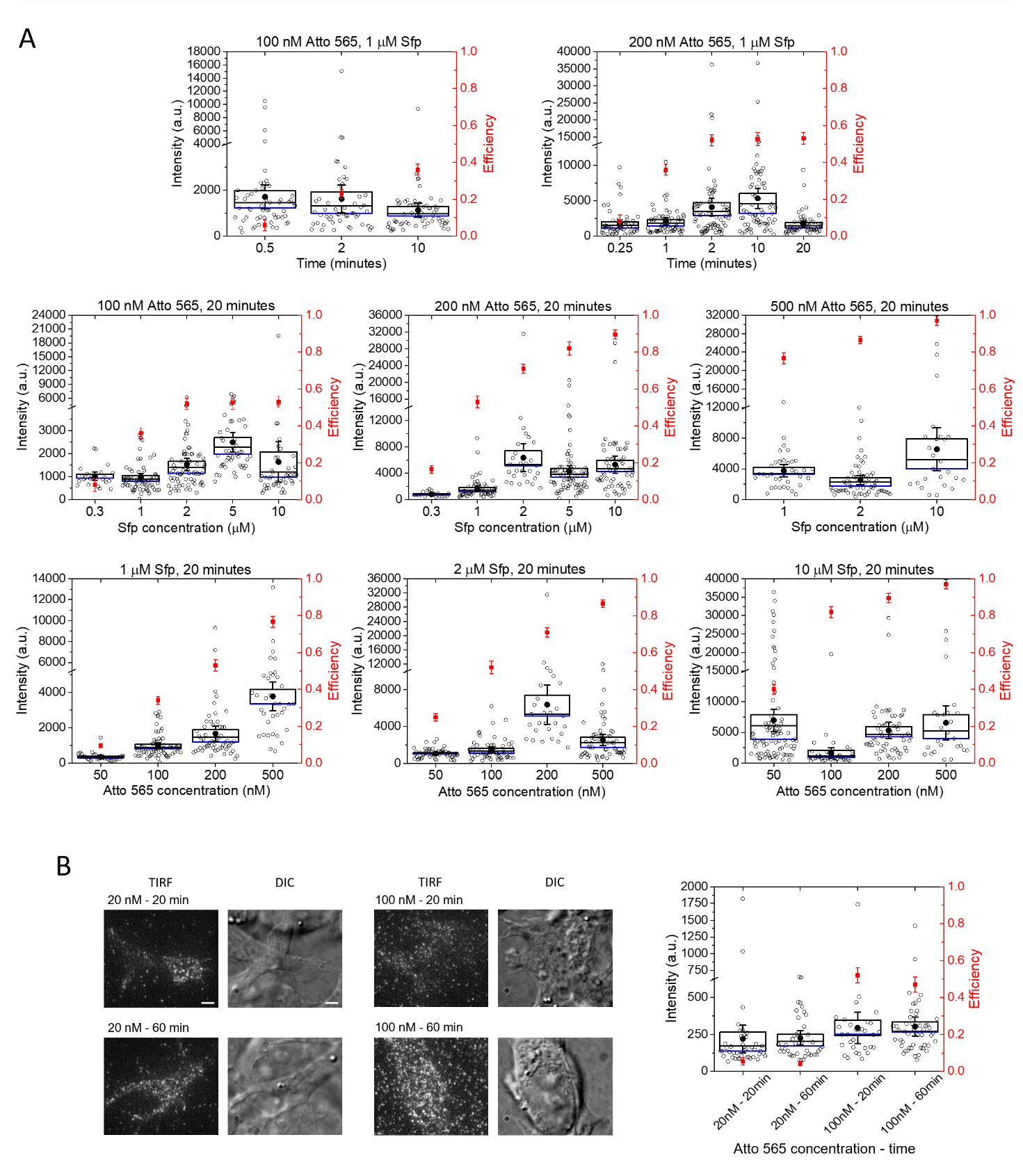

**Supplementary Fig.** **2** Density comparisons are not able to compare accurately labeling efficiencies. In black: mean intensities within labeled cells (background corrected, see Methods) are measured in different conditions in living (A) and fixed (B) cells. Boxes are mean ± SEM, whiskers are CI 95%, black dots are averages, blue lines are medians, empty circles are individual data from two independent replicates. The efficiencies measured with our method are reported in red (mean ± estimated SEM) for direct comparison (right y-axis). On the left y-axis, changes in scales are adopted to visualize the tails of the distributions. In (B), on the left, representative TIRF and DIC images are shown (scale bar: 5 μm)

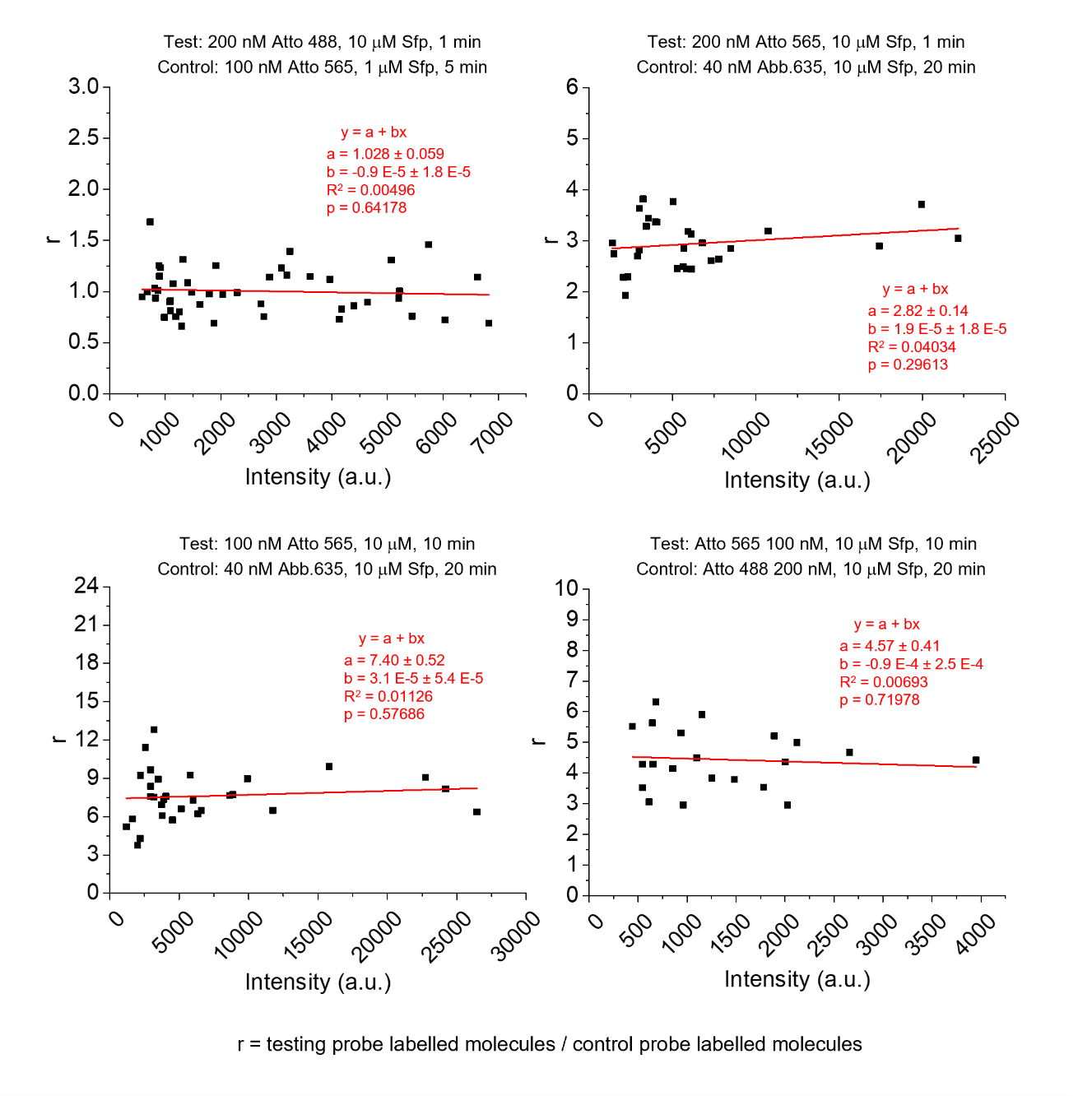

**Supplementary Fig. 3** Robustness of the method against expression variability. The plots show the ratio r between the numbers of receptors labeled with the test and the control probe (see Fig. 1), as a function of the labeled cell intensity detected in the channel of the test probe, used in the first labeling reaction. Different probes and reaction conditions are used, as stated for each plot. Spots are individual data from two independent replicates. In red, linear fitting curves with fit parameters are shown. p is the p-value for an F-test of whether the fitting model differs significantly from the model y=constant. In all cases, the slope is not significantly different from zero

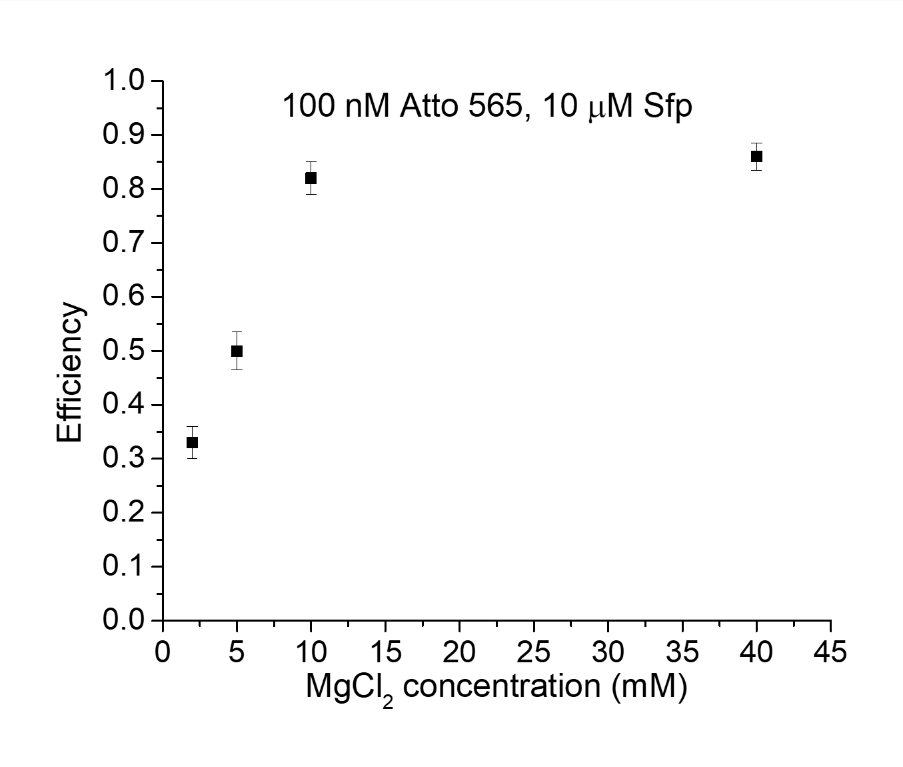

**Supplementary Fig. 4** Efficiency as a function of MgCl_2_ concentration. Measured efficiencies for Atto 565 (20-minutes reaction with indicated concentrations for dye and Sfp) using different MgCl_2_ concentrations in the labeling reaction. Data are reported as mean ± SEM estimate (see Methods) and are obtained from 12-32 analyzed cells in two independent replicates for each condition

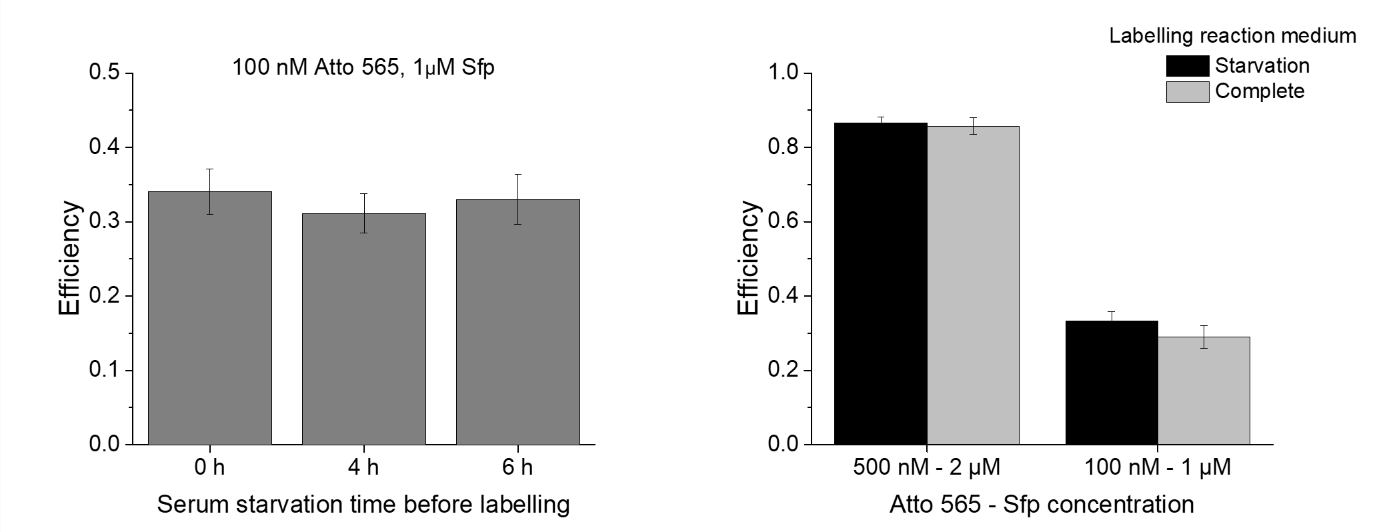

**Supplementary Fig. 5** Efficiency does not depend on serum starvation before and during labeling. (A) Measured efficiency for Atto 565 (20-minutes reaction with indicated concentrations) applying different serum starvation times before the reaction. (B) Efficiency comparison between labeling in starvation or complete medium (see Methods) at two different couple of dye-Sfp concentrations. Bars are mean ± SEM estimate (see Methods) and are obtained from 12-39 analyzed cells in each condition. Differences are not significant according to Anova (left graph) and Student (right graph) tests.

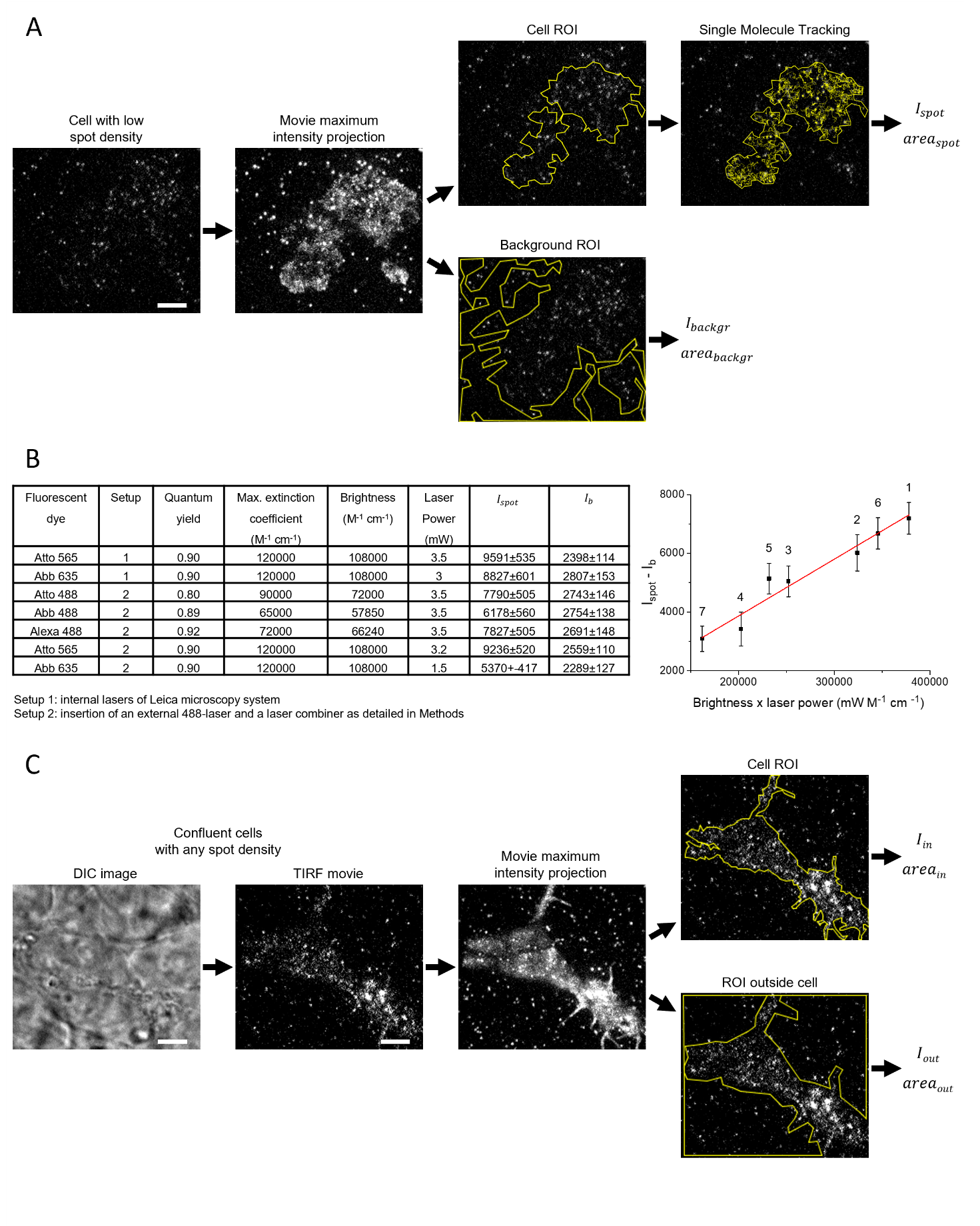

**Supplementary Fig. 6** Determination of the number of labeled receptors in the cell basal membrane. (A) Transfected cells with low moving spot densities are selected for TIRF movies acquisition (reported image is an example of a movie first frame). The movie maximum intensity projection is used to identify the contours of the basal membrane of transfected cells. Inside these contours (cell ROI), an analysis of single-particle tracking is performed to estimate the intensity and area of single labeled receptors ($I_{spot},{area}_{spot}$). An area without spots is selected outside the identified contours (background ROI), and $I_{backgr}$ and ${area}_{backgr}$ are estimated (see Methods). (B) Table reporting theoretical absorption and emission parameters, experimental setup and quantities measured with the procedure in panel A for the different fluorescent dyes used in the study. $I_{b}=\frac{I_{backgr}}{{area}_{backgr}}\times{area}_{spot}$ (see Methods). The graph on the right shows the direct proportionality between the theoretical dye brightness multiplied by the used laser power and the estimated spot intensity (background corrected). In black: experimental measures, mean ± SEM as in the table; numerical label of each measure indicates the corresponding row in the table; red: fitting curve y=ax. The data do not take into account the possibly different fraction of photons emitted within the bandwidth of the emission filters for dyes with different emission spectra, even if excited with the same laser, nor the fact that the excitation is not necessarily at the maximum of the excitation spectra. (C) Estimation of the number of labeled receptors. Samples with confluent cells are used (as seen in DIC mode); TIRF movies are acquired for transfected cells with any labeled spot density. Cell contours are identified like in panel A and integral intensity and area inside and outside these contours are evaluated. The extracted quantities are used in equation (5) together with the ones estimated in the step represented in panel A to obtain the number of labeled receptors. This number is then used to estimate the ratio of labeled receptors between two different channels in order to apply equations (1-4). Scale bars: 5 μm

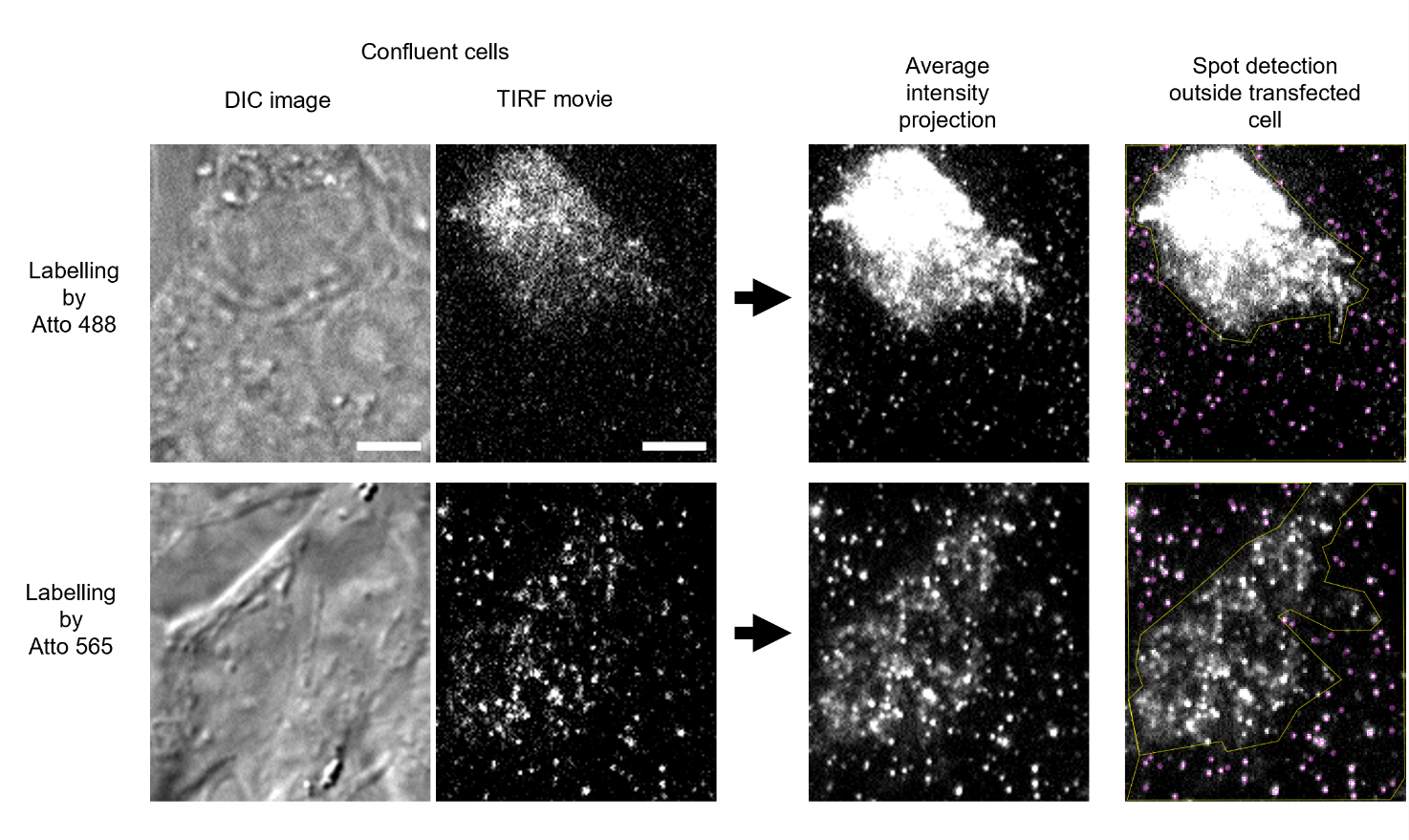

**Supplementary Fig. 7** Estimation of non-specific adhesion. The figure shows the steps performed for the analysis of non-specific adhesion. Samples are prepared with cells at full confluency. DIC image and the first image of the acquired TIRF movie are shown (scale bar: 5 μm). An average of the first 10 frames is used to improve the signal-to-noise ratio for single spot detection, performed outside the transfected cells (see Methods). In the last image of each row, the magenta circles represent spots detected with TrackMate plugin. First images row: example of a sample incubated with a reaction mix including Atto 488 dye; second images row: example of a sample incubated with a reaction mix including Atto 565 dye

| **Probe A: conditions** | **Probe B: conditions** | **Figure** |
| --- | --- | --- |
| Atto 565: 100 nM CoA-dye, 2 μM Sfp, 20 min | Abb. 635: 40 nM CoA-dye, 10 μM Sfp, 20 min | 2 |
| Atto 565: 100 nM CoA-dye, 10 μM Sfp, 10 min | Abb. 635: 40 nM CoA-dye, 10 μM Sfp, 20 min | 2 |
| Atto 565: 100 nM CoA-dye, 2 μM Sfp, 20 min | Abb. 635: 60 nM CoA-dye, 10 μM Sfp, 20 min | 6 |
| Atto 565: 200 nM CoA-dye, 5 μM Sfp, 20 min | Abb. 635: 100 nM CoA-dye, 1 μM Sfp, 20 min | 5B |

**Supplementary Table 1** Performed experiments of the type illustrated in Fig. 1A. The two labeling reactions described in each row are carried out in both sequential orders. The table reports used conditions for the two dyes and the figures including the obtained results

| **Test probe: conditions** | **Control probe: conditions** | **Reaction order** | **Figure** |
| --- | --- | --- | --- |
| Atto 565: all conditions in Fig. 2D | Abb. 635: 40 nM CoA-dye, 10 μM Sfp, 20 min | Test-control | 2D |
| Atto 565: 100 nM CoA-dye, 1 μM Sfp, 0.5, 1, 2, 5, 10, 20, 40 min | Abb. 635: 40 nM CoA-dye, 10 μM Sfp, 20 min | Test-control | 4A |
| Atto 565: 100 nM CoA-dye, 10 μM Sfp, 1, 2, 10, 20, 40 min | Abb. 635: 40 nM CoA-dye, 10 μM Sfp, 20 min | Test-control | 4A |
| Atto 565: 100 nM CoA-dye, 10 μM Sfp, 0.5 min | Atto 488: 200 nM CoA-dye, 10 μM Sfp, 5 min | Test-control | 4A |
| Atto 565: 100 nM CoA-dye, 10 μM Sfp, 5 min | Atto 488: 200 nM CoA-dye, 10 μM Sfp, 5 min | Control-test | 4A |
| Atto 565: 200 nM CoA-dye, 1 μM Sfp, 0.25, 0.5, 1, 2, 5, 10, 20 min | Abb. 635: 40 nM CoA-dye, 10 μM Sfp, 20 min | Test-control | 4A |
| Atto 565: 200 nM CoA-dye, 10 μM Sfp, 0.5, 1, 2, 5, 10, 20, 40 min | Abb. 635: 40 nM CoA-dye, 10 μM Sfp, 20 min | Test-control | 4A |
| Atto 565: 200 nM CoA-dye, 10 μM Sfp, 0.25 min | Atto 488: 200 nM CoA-dye, 10 μM Sfp, 5 min | Test-control | 4A |
| Atto 488: 500 nM CoA-dye, 5 μM Sfp, 20 min | Atto 565: 200 nM CoA-dye, 5 μM Sfp, 20 min | Test-control | 5C |
| Abberior 488: 500 nM CoA-dye, 5 μM Sfp, 20 min | Atto 565: 200 nM CoA-dye, 5 μM Sfp, 20 min | Test-control | 5C |
| Alexa 488: 500 nM CoA-dye, 5 μM Sfp, 20 min | Atto 565: 200 nM CoA-dye, 5 μM Sfp, 20 min | Test-control | 5C |
| Atto 488: 50 nM CoA-dye, 10 μM Sfp, 20 min | Atto 565: 100 nM CoA-dye, 1 μM Sfp, 5 min | Test-control | 5E |
| Atto 488: 100 nM CoA-dye, 10 μM Sfp, 20 min | Atto 565: 100 nM CoA-dye, 10 μM Sfp, 5 min | Test-control | 5E |
| Atto 488: 200 nM CoA-dye, 10 μM Sfp, 20 min | Atto 565: 100 nM CoA-dye, 10 μM Sfp, 5 min | Test-control | 5E |
| Atto 488: 500 nM CoA-dye, 10 μM Sfp, 20 min | Atto 565: 100 nM CoA-dye, 5 μM Sfp, 20 min | Test-control | 5E |
| Atto 488: 200 nM CoA-dye, 10 μM Sfp, 1, 2, 10, 40 min | Atto 565: 100 nM CoA-dye, 1 μM Sfp, 5 min | Test-control | 5E |
| Atto 488: 200 nM CoA-dye, 10 μM Sfp, 5 min | Atto 565: 100 nM CoA-dye, 10 μM Sfp, 20 min | Test-control | 5E |
| Atto 565 + 488: 65 + 130 nM CoA-dye, 20 μM Sfp, 20 min | Abb. 635: 40 nM CoA-dye, 10 μM Sfp, 20 min | Test-control | 6C |
| Atto 565 + 488: 65 + 130 nM CoA-dye, 20 μM Sfp, 20 min | Abb. 635: 40 nM CoA-dye, 10 μM Sfp, 20 min | Control-test | 6C |
| Atto 565 + 488: 100 + 200 nM CoA-dye, 20 μM Sfp, 2, 5, 10, 20 min | Abb. 635: 60 nM CoA-dye, 10 μM Sfp, 20 min | Test-control | 6C |
| Atto 565 + 488: 100 + 200 nM CoA-dye, 10 μM Sfp, 20 min | Abb. 635: 60 nM CoA-dye, 10 μM Sfp, 20 min | Test-control | 6C |
| Atto 565 + 488: 100 + 200 nM CoA-dye, 30 μM Sfp, 20 min | Abb. 635: 40 nM CoA-dye, 10 μM Sfp, 20 min | Test-control | 6C |
| Atto 565: 100 nM CoA-dye, 2 μM Sfp, 20 min | Abb. 635: 40 nM CoA-dye, 10 μM Sfp, 20 min | Test-control | Supplementary Fig. 1A |
| Atto 565: 100 nM CoA-dye, 2 μM Sfp, 20 min | Abb. 635: 40 nM CoA-dye, 10 μM Sfp, 20 min | Control-test | Supplementary Fig. 1A |
| Atto 565: 100 nM CoA-dye, 1 μM Sfp, 5 min | Abb. 635: 60 nM CoA-dye, 10 μM Sfp, 20 min | Test-control | Supplementary Fig. 1B |
| Atto 565: 100 nM CoA-dye, 1 μM Sfp, 5 min | Abb. 635: 40 nM CoA-dye, 10 μM Sfp, 20 min | Test-control | Supplementary Fig. 1B |
| Atto 565: 100 nM CoA-dye, 5 μM Sfp, 20 min | Abb. 635: 40 nM CoA-dye, 10 μM Sfp, 20 min | Test-control | Supplementary Fig. 1C |
| Atto 565: 100 nM CoA-dye, 5 μM Sfp, 20 min | Atto 488: 200 nM CoA-dye, 10 μM Sfp, 5 min | Control-test | Supplementary Fig. 1C |
| Atto 565: 100 nM CoA-dye, 10 μM Sfp, 10 min | Atto 488: 200 nM CoA-dye, 10 μM Sfp, 20 min | Test-control | Supplementary Fig. 1D |
| Atto 565: 100 nM CoA-dye, 10 μM Sfp, 10 min | Abb. 635: 40 nM CoA-dye, 10 μM Sfp, 20 min | Test-control | Supplementary Fig. 1D |
| Atto 565: 100 nM CoA-dye, 10 μM Sfp, 10 min | Abb. 635: 40 nM CoA-dye, 10 μM Sfp, 20 min | Control-test | Supplementary Fig. 1D |
| Atto 565: 100 nM CoA-dye, 10 μM Sfp, 20 min, 2, 5, 10, 40 mM MgCl_2_ | Abb. 635: 40 nM CoA-dye, 10 μM Sfp, 20 min | Test-control | Supplementary Fig. 4 |
| Atto 565: 100 nM CoA-dye, 1 μM Sfp, 20 min, 0,4,6 starvation hours | Abb. 635: 100 nM CoA-dye, 1 μM Sfp, 20 min | Test-control | Supplementary Fig. 5A |
| Atto 565: 500 nM CoA-dye, 2 μM Sfp, 20 min, starvation/No starvation | Abb. 635: 100 nM CoA-dye, 1 μM Sfp, 20 min | Test-control | Supplementary Fig. 5B |
| Atto 565: 100 nM CoA-dye, 1 μM Sfp, 20 min, starvation/No starvation | Abb. 635: 100 nM CoA-dye, 1 μM Sfp, 20 min | Test-control | Supplementary Fig. 5B |
| Atto 565: 20 nM CoA-dye, 2 μM Sfp, 20, 60 min | Abb. 635: 40 nM CoA-dye, 10 μM Sfp, 20 min | Test-control | Supplementary Fig. 2B |
| Atto 565: 100 nM CoA-dye, 2 μM Sfp, 20, 60 min | Abb. 635: 40 nM CoA-dye, 10 μM Sfp, 20 min | Test-control | Supplementary Fig. 2B |

**Supplementary Table 2** Performed experiments of the type illustrated in Fig. 1B. The table reports used conditions (dye: reaction parameters) for the test probe (first column) and the control probe (second column), the order of the reactions (third column), the figures including the obtained results (fourth column)

**Supplementary Video 1** Single-molecule imaging of labeled s6-TrkA receptors moving on the membrane of living cells. Labeling was performed with Atto 565 at the optimized conditions: 100 nM CoA-Atto 565, 10 μM Sfp, 10 mM MgCl_2_, 20 minutes. Scale bar: 5 μm. Related to Fig. 4C where a single image from this movie is shown

**Supplementary Video 2** Single-particle tracking analysis performed on Supplementary Video 1 with TrackMate plugin. In yellow: contours of the cell membrane and reconstructed trajectories. Scale bar: 5 μm. Related to Fig. 4C where an image from this analysis is shown

**Supplementary Video 3** Two-color single-molecule imaging of s6-TrkA receptors moving on the membrane of living cells and labeled simultaneously in two channels. Labeling was performed with 100 nM Atto 565 (right channel) + 200 nM Atto 488 (left channel), 20 μM Sfp, 10 mM MgCl_2,_ 5 minutes. Scale bar: 5 μm. Related to Fig. 6D where a single image from this movie is shown

**Supplementary Video 4** Single-particle tracking analysis performed on the Atto 565 channel of Supplementary Video 3 with the TrackMate plugin. In yellow: contours of the cell membrane and reconstructed trajectories. Scale bar: 5 μm. Related to Fig. 6D where an image from this analysis is shown

**Supplementary Video 5** Single-particle tracking analysis performed on the Atto 488 channel of Supplementary Video 3 with the TrackMate plugin. In yellow: contours of the cell membrane and reconstructed trajectories. Scale bar: 5 μm. Related to Fig. 6D where an image from this analysis is shown

**Supplementary Note 1**

As described in the main text (section: “Method to determine biomolecule-labeling efficiency" in the Results), the first labeling reaction on a cell with N labelable molecules labels $e_{A}\cdot N$ molecules, where $e_{A}$ is the probe efficiency. The second reaction (probe efficiency: $e_{B}$) labels $e_{B}\cdot\left( N - e_{A}N \right)$ on the same cell. The ratio between the number of molecules labeled in the first and in the second reaction is $r=\frac{e_{A}\cdot N}{e_{B}\cdot N\left( 1 - e_{A} \right)}$, where *N* can be simplified to obtain an expression independent on the total number of molecules. This allows having only two unknowns (the two efficiencies); inverting the order of labelling gives a second equation, as shown in the main text, and they can be solved for the two unknowns. It is important to note that the approach based on two dyes allows visualizing each cell in two detection channels at the same time, so that the number of molecules labeled after the first and after the second reaction are actually measured on each cell and *N* does not have any impact, being fixed for a fixed cell. This is a crucial point because typically different cells in a population (even a homogeneous one, like in a single petri dish) show gene expression variability due to different factors (from cell state and cell cycle to random phenomena, see also ref. 20 and 21 in the main text). Evaluation of the ratio r on each cell eliminates the impact of expression variability and allows therefore reliable estimations.

The starting idea of the method, i.e. the exploitation of two subsequent labeling reactions, could be applied in principle using a single detection channel with a single dye having efficiency *e*. In this case the ratio between the number of labeled molecules in a cell with N labelable molecules after the first and after the second reaction would be $r=\frac{Ne}{N(1+e-e^{2})}$, where the denominator includes molecules labeled in the first and in the second reaction. However, in this case, “N” usually is not strictly the same in the two measures, so its “simplification” is less straightforward. Indeed, in this case the samples would be imaged once after the first and a second time after the second reaction, and it would be very improbable to find and measure the same cells in the two visualization stages. So, *r* cannot be measured for each cell, but one must measure a mean intensity after each reaction on two different sets of cells. However, as shown also in Supplementary Figures 2 and 3, the variability for this mean intensity is extremely wide, and therefore the uncertainty would be high. Moreover and maybe more importantly, for living cells the number of labelled and labelable molecules could change between the two measurements. While for the method with two fluorophores, the two reactions are made immediately one after the other and the two channels are measured at the same time, for a single fluorophore there is a longer time between the two reactions (the time for the first imaging session) and a probably stress for the cells during the first step of imaging. So, also depending on the type of measurement (e.g. on whole cells, only on a compartment, only on the bottom membrane, etc.), there could be a sort of “recycling” that can make more labelable molecules available and remove some of the previously labelled ones, either because physically removed from the observation volume or because “destroyed”, including bleaching of the dye (that could happen also for non-living cells). Consequently, *N* is not constant between the two sets of cells (it can change, at least it is a statistic), it cannot be simply simplified in the equations, and the determination of the labeling efficiency is less precise than with two different dyes.

However, application with a single dye is a feasible simplified way to obtain approximate estimations in some cases, e.g., when a single reactive dye is available; another simplification is that, using a single dye, one can measure fluorescence intensities without the need for converting them in number of molecules. Indeed, by using fixed dye and consequently fixed microscope settings (such as detection channel), the intensity is proportional to the number of molecules with a proportionality constant that is equal for numerator and denominator in the expression of *r*.

**Supplementary Note 2**

In experiments of the kind represented in fig. 1A, with reactions performed in both orders, we measure the ratios r and r' for each analyzed cell; then, the values averaged on all cells are inserted in equations (1) and (2) to calculate the efficiencies.

The uncertainty for $e_{A}$ calculated from equation (1) is $\sigma_{e_{A}}=\frac{1}{r^{'}(r+1)}\sqrt{\frac{\left( r^{'}+1 \right)^{2}}{{(r+1)}^{2}}\sigma_{r}^{2}+\frac{\sigma_{r^{'}}^{2}}{{r^{'}}^{2}}}$ , and the uncertainty for $e_{B}$ calculated from equation (2) is $\sigma_{e_{B}}= \frac{1}{r(r^{'}+1)}\sqrt{\frac{\sigma_{r}^{2}}{r^{2}}+\frac{{(r+1)}^{2}}{\left( r^{'}+1 \right)^{2}}\sigma_{r^{'}}^{2}}$, where$\sigma_{r}$ and $\sigma_{r^{'}}$ are respectively the SEM of the ratios $r$ and $r^{'}$, measured in repeated and independent experiments.

In experiments of the kind represented in fig. 1B, we measure the ratio r for each analyzed cell; then, the value averaged on all cells is inserted in equation (3) or (4) to calculate the efficiency, together with the mean value of the control probe efficiency previously measured in different and independent experiments.

The uncertainty for $e_{A}$ calculated from equation (3) is $\frac{1}{{(1+re_{B})}^{2}}\sqrt{\sigma_{r}^{2}+r^{2}\sigma_{e_{B}}^{2}}$, and the uncertainty for $e_{B}$ calculated from equation (4) is $\frac{1}{r(e_{A}-1)}\sqrt{\frac{e_{A}^{2}\sigma_{r}^{2}}{r^{2}}+\frac{\sigma_{e_{A}}^{2}}{{(e_{A}-1)}^{2}}}$, where$\sigma_{r}$ and $\sigma_{e_{B}}$ (or $\sigma_{e_{A}}$) are respectively the SEM of the ratio $r$ just measured and the SEM of the control probe efficiency $e_{B}$ (or $e_{A}$) previously determined.

All reported formula for uncertainties were calculated using propagation error for independent measurements. Since efficiencies ($y$, below) are calculated from the results $x_{i}$ of various independent experiments with different sample numbers ($y=f\left( x_{1},x_{2},\ldots\right)$), the effective degrees of freedom $\nu_{eff}$ that have to be used for comparison statistical tests have been calculated via the Welch-Satterthwaite approximation:

$$\nu_{eff}\left( y \right)=\frac{u_{c}^{4}(y)}{\sum_{i=1}^{N} \frac{u_{i}^{4}(y)}{\nu_{i}}}$$

where$u_{c}^{2}\left( y \right)=\sum_{i=1}^{N} c_{i}^{2}u^{2}(x_{i})$ is the combined variance associated with the output estimate $y$ ($c_{i}=\frac{\partial f}{\partial x_{i}}$ in the case of propagated uncertainty); $u_{i}^{2}\left( y \right)=\left[ c_{i} u(x_{i}) \right]^{2}$ is the component of the combined variance $u_{c}^{2}(y)$ generated by the estimated variance $u^{2}(x_{i})$ of the input estimate $x_{i}$, and $\nu_{i}$ are the degrees of freedom of $u(x_{i})$ (e.g., $\nu_{i}=n_{i}-1$ if $x_{i}$ is an average of $n_{i}$ data).

Finally, using (3) or (4) to calculate the test efficiency $e_{test}$ from the ratio *r* and the control efficiency $e_{ctrl}$, the effective degrees of freedom are:

$$\nu_{eff}\left( e_{test} \right)=\frac{u_{c}^{4}(e_{test})}{\frac{1}{\nu_{r}}\left( \frac{\partial e_{test}}{\partial r}u(r) \right)^{4}+\frac{1}{\nu_{e_{ctrl}}}\left( \frac{\partial e_{test}}{\partial e_{ctrl}}u(e_{ctrl}) \right)^{4}}$$

If $x_{i}$ is itself obtained indirectly from other measurements, $\nu_{i}$ is the effective degree of freedom calculated as written above. This is the case for $e_{ctrl}$, obtained from previous measurements.

**Supplementary Note 3**

When making an experiment where a homo-interaction is studied by labeling the same kind of molecule alternatively with two different fluorophores, one usually measures something regarding the couples of molecules labelled with different fluorophores (e.g. FRET, or number of couples), and therefore wants to maximize their number. Considering, for simplicity, a case in which all molecules are labeled with one of two fluorophores B or G, and where all the molecules are in a couple, there will be a fraction x of molecules labeled with B and (1-*x*) with G. If the number of molecules is high enough, if the first molecule in couple is labelled with B (probability x), the other one could be B with probability *x* and G with probability (1-*x*) (independent probabilities). The same applies if the first molecule is G (probability 1-*x*). Therefore, one will find a couple BB with probability *x*^2^, GG with probability (1-*x*)^2^, and BG (without considering the order, since BG and GB are undistinguishable) with probability 2*x*(1-*x*) (note that the sum of the three probabilities gives 1, as it should be). This last probability has a maximum equal to ½ for *x*=½, therefore it is desirable to have the same labeling efficiency in the two channels.
